# VEGFA mRNA-LNP promotes biliary epithelial cell-to-hepatocyte conversion in acute and chronic liver diseases and reverses steatosis and fibrosis

**DOI:** 10.1101/2023.04.17.537186

**Authors:** Fatima Rizvi, Yu-Ri Lee, Ricardo Diaz-Aragon, Juhoon So, Rodrigo M. Florentino, Anna R. Smith, Elissa Everton, Alina Ostrowska, Kyounghwa Jung, Ying Tam, Hiromi Muramatsu, Norbert Pardi, Drew Weissman, Alejandro Soto-Gutierrez, Donghun Shin, Valerie Gouon-Evans

## Abstract

The liver is known for its remarkable regenerative ability through proliferation of hepatocytes. Yet, during chronic injury or severe hepatocyte death, proliferation of hepatocytes is exhausted. To overcome this hurdle, we propose vascular-endothelial-growth-factor A (VEGFA) as a therapeutic means to accelerate biliary epithelial cell (BEC)-to-hepatocyte conversion. Investigation in zebrafish establishes that blocking VEGF receptors abrogates BEC-driven liver repair, while VEGFA overexpression promotes it. Delivery of VEGFA via non-integrative and safe nucleoside-modified mRNA encapsulated into lipid-nanoparticles (mRNA-LNP) in acutely or chronically injured mouse livers induces robust BEC-to-hepatocyte conversion and reversion of steatosis and fibrosis. In human and murine diseased livers, we further identified VEGFA-receptor KDR-expressing BECs associated with KDR-expressing cell-derived hepatocytes. This defines KDR-expressing cells, most likely being BECs, as facultative progenitors. This study reveals novel therapeutic benefits of VEGFA delivered via nucleoside-modified mRNA-LNP, whose safety is widely validated with COVID-19 vaccines, for harnessing BEC-driven repair to potentially treat liver diseases.

**Graphical Abstract:** 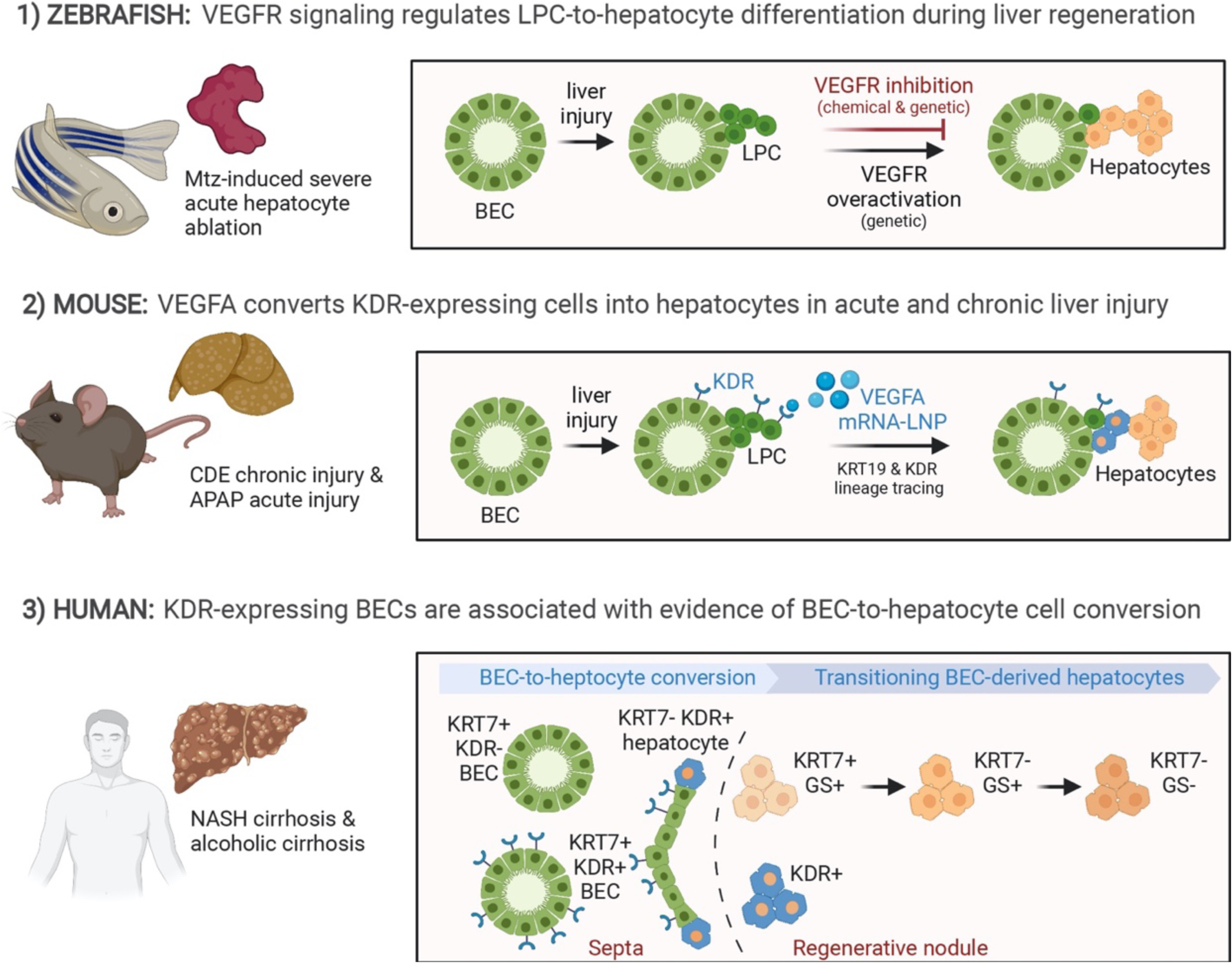

**Highlights:**

1. Complementary mouse and zebrafish models of liver injury demonstrate the therapeutic impact of VEGFA-KDR axis activation to harness BEC-driven liver regeneration.
2. VEGFA mRNA LNPs restore two key features of the chronic liver disease in humans such as steatosis and fibrosis.
3. Identification in human cirrhotic ESLD livers of KDR-expressing BECs adjacent to clusters of KDR+ hepatocytes suggesting their BEC origin.
4. KDR-expressing BECs may represent facultative adult progenitor cells, a unique BEC population that has yet been uncovered.

## Introduction

Mortality related to end stage liver disease (ESLD) is ranked as the 12^th^ most common cause of death in the US. Liver transplantation remains the only treatment for ESLD, but this procedure is critically challenged by the shortage of organ donors. The remarkable ability of the liver to regenerate by proliferation of mature hepatocytes is well documented (Michalopoulos, 2017), yet in the case of severe acute hepatocyte death or chronic ESLD, proliferation of mature cells becomes exhausted (Duncan et al., 2009; Stanger, 2015) due to the escalating progression of steatosis, inflammation, fibrosis and irreversible cirrhosis. In these cases, the presence of alternative precursors of hepatocytes that derive from biliary epithelial cells (BECs) has been postulated, and these cells have been referred to by various names, including liver progenitor cells (LPCs) (Rodrigo-Torres et al., 2014). Expansion of LPCs from the BEC compartment, a process described as ductular reaction (DR), is present in virtually all chronic and acute human liver diseases, suggesting an alternative regeneration process by which BECs proliferate and differentiate into hepatocytes (Boulter et al., 2013; Gouw et al., 2011; Lowes et al., 1999; Popper et al., 1957; Roskams and Desmet, 1998; Roskams et al., 2003; Roskams et al., 2004; Sato et al., 2019; Shin and Kaestner, 2014; Turanyi et al., 2010; Van Haele et al., 2019), the BEC-driven liver regeneration. The evidence for the contribution of BECs to *de novo* hepatocytes in humans is illustrated by the presence of hepatocytes expressing the BEC marker EpCAM emerging from highly proliferative DR areas in advanced cirrhotic livers (Yoon et al., 2011), detection of “bi-phenotypic cells” (Deng et al., 2018) or “ductular hepatocytes” (Haque et al., 1996) positive for both the BEC marker KRT19 and the hepatocyte marker HNF4α (Deng *et al*., 2018) or HepPar1 (Haque et al., 1996), and observation of budding of hepatocyte-like cells expressing the hepatocytic marker glutamine synthetase (GS) from BECs within the DRs (Falkowski et al., 2003; Fleming and Wanless, 2013; Hytiroglou and Theise, 2018; Stueck and Wanless, 2015; Wanless et al., 2000). Specifically, Stueck and colleagues have quantified the emerging immature hepatocytes budding from KRT19+ BECs within the septa developed in human cirrhotic livers, and estimated that they represented up to 70% of hepatocytes (Stueck and Wanless, 2015). Importantly, a recent study showed that aberrant GS positivity adjacent to portal tracts is significantly associated with regressed cirrhosis in humans (Hadi et al., 2020), suggesting the clinical benefit of LPC-derived hepatocytes in resolving human cirrhosis. In an attempt to lineage trace the LPCs among the DRs in human cirrhotic livers, Lin and colleagues have used mutational analysis in mitochondrial DNA encoding cytochrome c oxidase enzyme, and showed the descent of hepatocytes within monoclonal regenerative nodules from adjacent LPC-associated DRs (Lin et al., 2010). In mice, lineage tracing studies have confirmed that the first and massive response to liver injury is the proliferation of mature hepatocytes (Malato et al., 2011; Overturf et al., 1997; Schaub et al., 2014; Yanger et al., 2014). Yet, consistent with an alternative BEC-mediated liver repair identified in human liver diseases, the BEC origin of *de novo* hepatocytes has been demonstrated in mouse models in which hepatocyte proliferation was significantly impaired by lack of Mdm2 (Lu et al., 2015), deficiency in β1 integrin (Raven et al., 2017) or β-catenin (Russell origin of the majority of hepatocytes after near complete ablation of hepatocytes (Choi et al., 2014; He et al., 2014).

This emerging literature raises the exciting possibility that BECs could be harnessed for therapeutic purposes to produce *de novo* hepatocytes and restore liver function. The current limitation for the efficient therapeutic use of BECs is the lack of druggable pathways that efficiently trigger the BEC-to-hepatocyte conversion. In this study, we propose that VEGFA promotes BEC-to-hepatocyte conversion and rescues liver function using complementary mouse and zebrafish liver injury models. Although VEGFA signaling is a known master regulator of angiogenesis, previous studies provide evidence for its role in improving liver regeneration after injury in rodents (Bockhorn et al., 2007; Ding et al., 2010; LeCouter et al., 2003; Yang et al., 2014). The last two studies indicated that VEGFA binds KDR expressed on liver sinusoidal endothelial cells that in turn secrete hepatocyte mitogens, thereby promoting the proliferation of hepatocytes. The role of VEGFA in accelerating BEC-driven regeneration was suggested in a rat injury model in which hepatocyte proliferation was compromised (Oe et al., 2005); however, it is not known yet if VEGFA promotes BEC-to-hepatocyte conversion. A direct effect of VEGFA on the BEC lineage is possible as KDR is expressed in BECs following liver injury in rats (Gaudio et al., 2006), in developing BECs generated from human induced pluripotent stem cells (Dianat et al., 2014), and in ductal plates in the developing human fetal biliary system (Fabris et al., 2006). Interestingly, our previous studies using embryonic stem cell differentiation cultures identified a liver progenitor cell expressing KDR whose differentiation into hepatocyte-like cells was inhibited when KDR signaling was chemically abrogated (Goldman et al., 2013). These data suggested a novel role for the ligand VEGFA to promote hepatocyte fate decision from a KDR-expressing precursor. Here, our complementary zebrafish and mouse lineage tracing models supported by analyses of human end stage liver disease (ESLD) specimens reveal that VEGFA, delivered with the clinically safe and non-integrative lipid nanoparticle-encapsulated nucleoside-modified mRNA (mRNA-LNP) platform (Pardi et al., 2015), induces BEC-to-hepatocyte differentiation through cell conversion of KDR-expressing BECs acting as facultative liver progenitor cells. Importantly, this study demonstrates the therapeutic benefit of VEGFA mRNA-LNP to harness BEC-driven liver regeneration and resolve liver diseases with potential clinical applications. This study uncovers a novel application of mRNA-LNP for tissue regeneration that departs from its original applications as protein replacement or immunization.

## Results

### VEGFR signaling regulates BEC-to-hepatocyte conversion during liver regeneration in zebrafish

To investigate the molecular mechanisms underlying BEC-driven liver regeneration, we had previously performed a chemical screen (Ko et al., 2016) using the zebrafish hepatocyte ablation model, *Tg(fabp10a:CFP-NTR)^s931^*, in which metronidazole (Mtz) treatment specifically ablates virtually all nitroreductase (NTR)-expressing *fabp10a^+^* hepatocytes, thereby eliciting BEC-driven liver regeneration (Choi *et al*., 2014^)^. Through this screen, we reported that suppressing VEGFR signaling with two VEGFR inhibitors, SU5416 (Fong et al., 1999) and SU4312 (Sun et al., 1998), significantly reduced the size of regenerating livers but did not affect the induction of early hepatocyte markers Prox1 and HNF4α in BEC-derived liver progenitor cells (LPCs) (Ko *et al*., 2016), suggesting normal BEC-to-LPC dedifferentiation. We here sought to determine whether VEGFR signaling regulates the next step in BEC-driven liver regeneration, LPC-to-hepatocyte differentiation. To distinguish BEC-derived hepatocytes, we used a Notch zebrafish reporter line, Tg(Tp1:H2B-mCherry)s939, which expresses histone 2B (H2B) and mCherry fusion proteins in BECs (Choi *et al*., 2014). A long half-life of H2B-mCherry allows for tracing the lineage of BECs; thus, BECs are strong H2B-mCherry+ and BEC-derived hepatocytes are weak H2B-mCherry+ (Choi *et al*., 2014). Treating *Tg(fabp10a:CFP-NTR)* larvae with SU5416 from ablation 18 hours (A18h) to regeneration 24 hours (R24h) significantly reduced the size of regenerating livers (Figure 1A), as reported (Ko *et al*., 2016). Importantly, the SU5416 treatment greatly suppressed the expression of the hepatocyte marker Bhmt at R24h (Figure 1A), suggesting a defect in LPC-to-hepatocyte differentiation. This defect was confirmed with two additional hepatocyte markers, *gc* and *f5* (Figure 1B). Complimentary to the pharmacological inhibition, we genetically blocked VEGFR signaling using the Tg(hs:sflt1)bns80 line, which expresses a soluble form of VEGFR1 (sFlt1), a decoy receptor for VEGFA, VEGFB, and PlGF (Gaudio *et al*., 2006), upon heat-shock (Matsuoka et al., 2017). sFlt1 overexpression also reduced Bhmt expression and liver size in regenerating livers at R26h (Figure 1C), phenocopying the effects of the SU5416 treatment.

**Figure 1.**
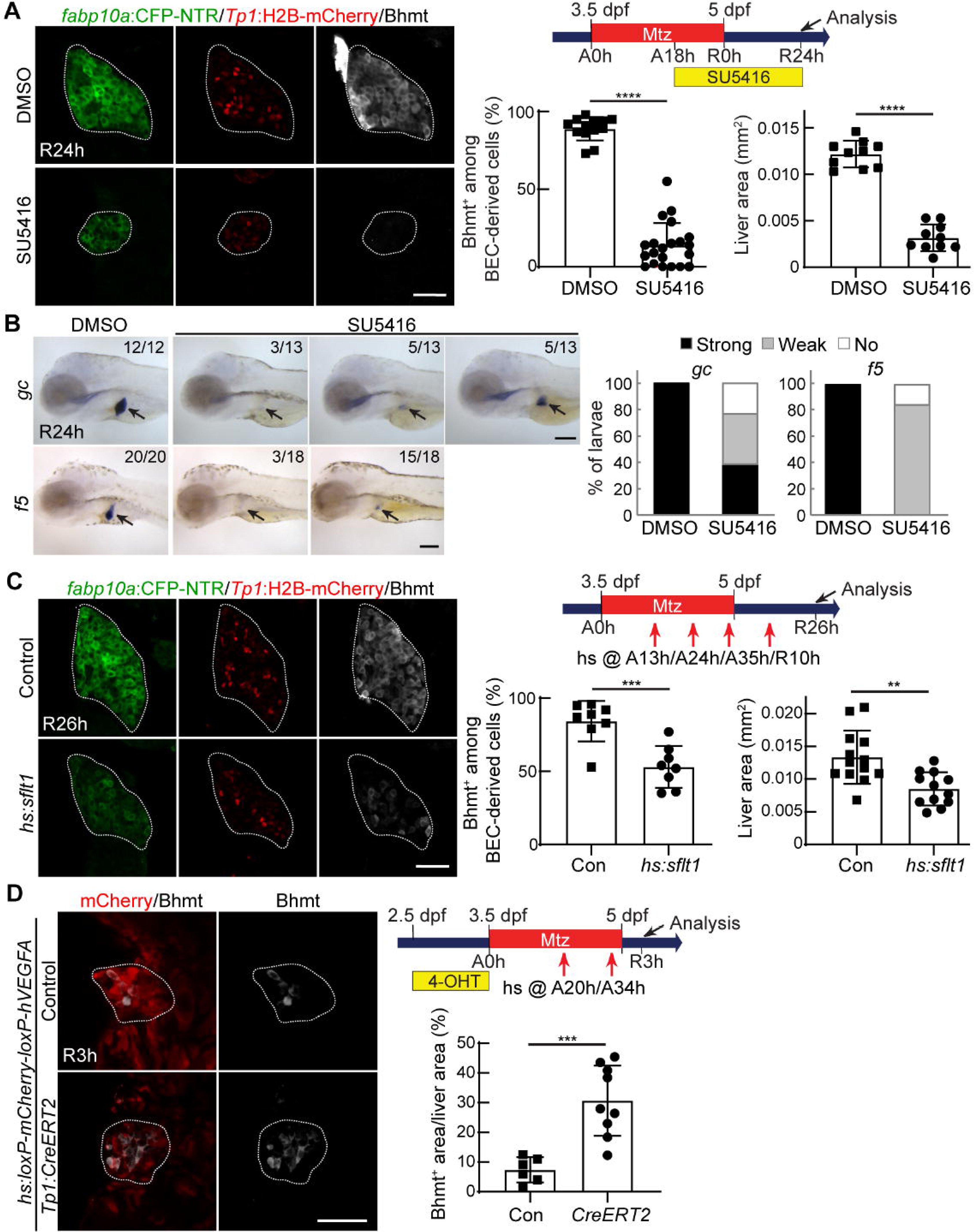
VEGFR signaling regulates BEC-driven liver regeneration in zebrafish. **(A)** Single-optical section images showing the expression of Bhmt (gray), *Tp1:*H2B-mCherry (red), and *fabp10a:*CFP-NTR (green) in regenerating livers (dotted lines) at R24h. Scheme illustrates the periods of Mtz and SU5416 treatments and an analysis stage. Quantification of the percentage of hepatocytes (Bhmt^+^) among BEC-derived cells (H2B-mCherry^+^) and quantification of liver size are shown. **(B)** Whole-mount in situ hybridization images showing *gc* and *f5* expression (arrows) in regenerating livers at R24h. Numbers in the upper-right corner indicate the proportion of larvae exhibiting the phenotype shown. Based on the levels of hepatic *gc* and *f5* expression, larvae were divided into three groups: no, weak, and strong. **(C)** Single-optical section images showing the expression of *Tp1:*H2B-mCherry, *fabp10a:*CFP-NTR, and Bhmt in regenerating livers (dashed lines) at R26h. To overexpress sFlt1, *Tg(hs:sflt1)* larvae were heat-shocked four times at A13h, A24h, A35h, and R10h. Quantification of the percentage of Bhmt^+^ among BEC-derived cells and quantification of liver size are shown. **(D)** Maximum projection images showing the expression of *hs:loxP*-mCherry-*loxP* and Bhmt in regenerating livers (dashed lines) at R3h. The *Tg(Tp1:CreERT2)* and *Tg(hs:loxP-mCherry-loxP-hVEGFA)* lines were used to express hVEGFA in a subset of BEC-derived cells during regeneration. Larvae were treated with 10 µM 4-OHT from 2.5 to 3.5 dpf for 24 hours, heat-shocked twice at A20h and A34h, and harvested at R3h. Quantification of the percentage of Bhmt^+^ area in the liver is shown. Data are presented as mean ± SEM. **p<0.01, ***p<0.001, and ****p<0.0001; statistical significance was calculated using an unpaired two-tailed t-test. Scale bars: 50 (A,C,D), 100 (B) µm.

Inversely, we next investigated if the overactivation of VEGFR signaling could promote LPC-to-hepatocyte differentiation. We used the *Tg(hs:loxP-mCherry-loxP-hVEGFA)^mn32^* line, which expresses human VEGFA165 upon heat-shock following Cre-mediated, mCherry-loxP excision (Hoeppner et al., 2012b), together with the Tg(Tp1:CreERT2)s959 line that expresses CreERT2 in BECs. Larvae were treated with 4-OHT from 2.5 to 3.5 days post-fertilization (dpf) to induce the Cre-mediated excision of mCherry-loxP in BECs, heat-shocked twice at A20h and A34h to trigger expression of VEGFA165, and harvested at R3h. hVEGFA expression in a subset of BEC-derived cells (BECs and LPCs) increased Bhmt expression in regenerating livers at R3h compared with controls (Figure 1D), indicating that VEGFR overactivation is sufficient to promote LPC-to-hepatocyte differentiation. Altogether, the zebrafish data demonstrate the essential role of VEGFR signaling in BEC-driven liver regeneration, particularly in LPC-to-hepatocyte differentiation.

### VEGFA mRNA-LNP induces BEC-to-hepatocyte conversion in CDE diet-induced chronic liver injury in mice and resolves liver damage

Given the impact of VEGFR activation in promoting BEC-driven liver regeneration in zebrafish, we interrogated its clinical benefit in restoring liver function in a chronic liver injury model in mice. To trace hepatocytic fate of BECs, liver injury was induced in *Krt19*-Cre^ERT^, *R26*^LSL^ tdTomato^39^ (Raven *et al*., 2017) mice with the choline-deficient diet supplemented with 0.1% ethionine (CDE) following a single injection of AAV8-*Tbg*-p21 (CDE/p21 model) (Raven *et al*., 2017) (Figure 2A), which recapitulates some key features of the human non-alcoholic steatohepatitis (NASH) liver disease including steatosis, fibrosis, hepatocyte senescence, and invasive ductular reaction (Raven *et al*., 2017). Injection of AAV8-*Tbg*-p21 triggered p21 expression in hepatocytes and was used to induce hepatocyte senescence (Raven *et al*., 2017; Russell *et al*., 2018), a common feature seen in human chronic liver disease (Marshall et al., 2005; Richardson et al., 2007). We chose to deliver VEGFA in mice via injection of nucleoside-modified mRNA-LNP allowing controlled transient expression of VEGFA in the liver for better clinical translation. We have recently demonstrated efficacy of mRNA-LNP as a tool to transiently express regenerative factors in the liver (Everton et al., 2021; Rizvi et al., 2021) and that their translation into proteins can last a few days, enough time to revert some features of liver diseases (Everton *et al*., 2021; Rizvi *et al*., 2021). For the present study, robust levels of VEGFA proteins in the serum were detected as early as 5 hours after one intravenous injection of 10μg/20g body weight of VEGFA mRNA-LNPs (Figure 2B). The levels remained high for the following 24 hours, yet rapidly decreased to baseline levels 48 hours later and were no longer detected by 72 hours. Mice were administered two injections of either VEGFA mRNA-LNP or control LNP (formulated with untranslatable Poly(C) RNA or neutral firefly luciferase-encoding mRNA) after the diet was over. Strikingly, large patches of diffused tdTomato+ clusters appeared in all liver lobes of VEGFA mRNA-LNP-treated mice, representing mostly hepatocytes (Figure 2C, yellow outline, Figure 2D). In contrast, in control LNP-treated mice, tdTomato+ clusters were sporadic and small (Figure 2C, Figure 2D), with the vast majority composed of BECs (Figure 2D, see numerous * areas). tdTomato+ hepatocytes (Figure 2d, arrowheads) were adjacent to tdTomato+ BECs (Figure 2D, arrows), supporting their BEC origin. Functionally, the tdTomato+ hepatocytes were equally efficient in storing glycogen as the adjacent tdTomato-hepatocytes (Figure 2E). Quantification of tdTomato+ hepatocytes by flow cytometry confirmed that VEGFA mRNA-LNPs significantly augment BEC-to-hepatocyte conversion (Figure 2F). Since the lineage tracing efficiency varies greatly among *Krt19*-Cre^ERT^, *R26*^LSL^ tdTomato mice, the percent labeled BEC population estimated from the non-parenchymal fraction (Figure S1A) was used to adjust for lineage tracing discrepancy and define the adjusted % tdTomato+ hepatocytes for each mouse (Figure 2F). We thus estimated that VEGFA mRNA-LNPs promote 5.3-fold greater % tdTomato+ hepatocytes as compared to control LNP-treated mice. In contrast to the reported role of VEGFA in promoting BEC proliferation (Gaudio *et al*., 2006), here the extent of DR was similar between the two groups (Figure S1B), most likely due to greater BEC-to-hepatocyte conversion in VEGFA treated group, as supported by lower density of KRT7+ BECs in areas associated with tdTomato+ hepatocytes (Figure S1C). Importantly, for clinical translation of VEGFA mRNA-LNP, VEGFA-mediated conversion of BECs to hepatocytes was consistently associated with complete reversion of fibrosis and steatosis (Figure 2G, 2H) as quantified with trichrome and LipidSpot staining, respectively. In line with these findings, the serum levels of cholesterol in the VEGFA mRNA-LNP-treated group were significantly higher, validating increased lipid clearance from hepatocytes as compared to control LNP-treated group (Figure 2I). Overall, treatment of the CDE/p21 mouse model with two injections of VEGFA mRNA-LNP fully reverts steatosis and fibrosis, the two key features observed in NASH patients, suggesting a potential clinical benefit of VEGFA mRNA-LNP to alleviate the human NASH disease.

**Figure 2.**
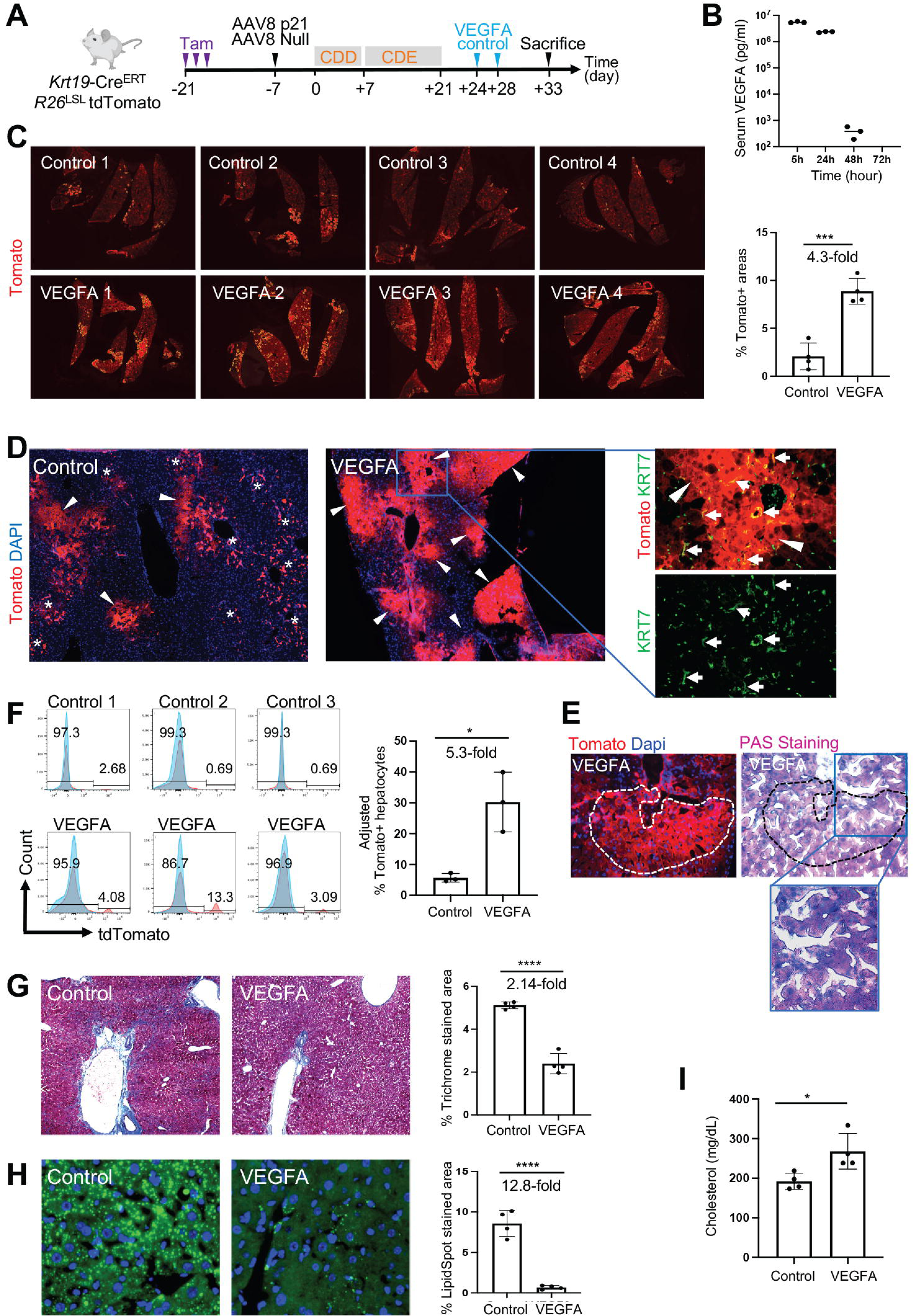
VEGFA mRNA-LNP administration induces BEC-to-hepatocyte conversion and promotes liver repair in CDE/p21-induced chronic liver injury in mice. **(A)** Scheme showing key interventions in the experimental design using the *KRT19*-Cre^ERT^, *R26*^LSL^tdTomato mice. AAV8-Tbg-p21 vector was administered to induce hepatocyte senescence. Injury was induced by CDE diet for 2 weeks, followed by VEGFA mRNA-LNP or control Poly(C) RNA-LNP injections, 10 μg/20g body weight. **(B)** Human-specific VEGFA ELISA in mouse serum (n=3), injected with VEGFA mRNA-LNPs. Serum was collected 5h, 24h, 48h, and 72h after injection. **(C)** Immunofluorescent images showing tdTomato+ clusters (outlined yellow) in livers of mice given two injections of either control Poly(C) RNA-LNP (n=4) or VEGFA mRNA-LNP (n=4). The bar graph shows quantification of tdTomato+ areas in the two groups. **(D)** Representative images of tdTomato+ (red) and KRT7+ (green) liver cells in Poly(C) RNA-LNP- or VEGFA mRNA-LNP-treated mice. The close-up images highlight tdTomato+ hepatocytes (arrowheads) adjacent to tdTomato+Krt7+ BECs (arrows) in VEGFA-mRNA-LNP-treated mice. * represents tdTomato+ BEC areas. **(E)** tdTomato and PAS staining on serial sections of liver tissue demonstrating glycogen storage in tdTomato+ hepatocytes. **(F)** Histograms from flow cytometry of hepatocyte fraction isolated from mouse livers. Values of red histograms represent % tdTomato+ population in the hepatocyte fraction. Blue histograms represent hepatocytes from a control non-tdTomato background run simultaneously with the experimental mice. Bar graph shows the total % tdTomato+ hepatocytes calculated by extrapolating the lineage tracing efficiency as 100% across all mice. **(G)** Representative brightfield images and corresponding bar graph showing quantification of % trichrome stained area estimated from at least three different fields in each mouse in the two groups (n=4 per group). **(H)** LipidSpot staining showing the accumulation of lipid droplets (green) in hepatocytes. The bar graph shows quantification of % LipidSpot stained area averaged from three different fields in each mouse in the two groups (n=4 per group). **(I)** Bar graph depicting total serum cholesterol levels in mice. Numerical data are presented as mean ± s.d. P values were determined by two-tailed Student’s t-test, *p< 0.05, **p< 0.01, ***p < 0.001, ****p<0.0001.

Since VEGFA significantly promotes emergence of BEC-derived hepatocyte in the CDE/p21 mice, we questioned the significance of p21-induced hepatocyte senescence in this process. Indeed, it has been reported by many groups that BEC-driven liver repair occurs almost exclusively in mouse models in which hepatocyte proliferation was impaired (Lu *et al*., 2015; Raven *et al*., 2017; Russell *et al*., 2018). Therefore, we asked whether suppression of hepatocyte proliferation was a prerequisite for VEGFA-mediated BEC-to-hepatocyte conversion. We thus administered VEGFA mRNA-LNP in CDE-fed mice that were not pre-injected with AAV8-*Tbg*-p21 (Figure S1D). In both control-LNP- and VEGFA mRNA-LNP-treated groups, the DR was much lower than in AAV8-*Tbg*-p21 administered mice (compare Figure S1E with Figure S1B), indicating that hepatocyte senescence accelerates BEC expansion. Control-LNPs group did not generate any tdTomato+ hepatocytes as expected (Raven *et al*., 2017). However, we consistently observed emergence of tdTomato+ hepatocyte clusters in the VEGFA mRNA-LNP-treated group, although sparse, in a specific spatial pattern (Figure S1F). The clusters were always adjacent to hepatocytes that naturally induced endogenous p21 expression (Figure S1G, panel 1,2,3). These results demonstrate that p21-mediated suppression of hepatocyte proliferation not only promotes BEC proliferation but also their conversion to hepatocytes. Importantly, VEGFA mRNA-LNP in the presence of hepatocyte proliferation can still trigger BEC-to-hepatocyte conversion, yet combined with suppression of hepatocyte proliferation, potentializes cell conversion to a clinically relevant extent and reverses steatosis and fibrosis.

### VEGFA promotes BEC-to-hepatocyte conversion in APAP-induced acute liver injury in mice

Given that acute acetaminophen (APAP) toxicity induces ductular reaction in mice (Kofman et al., 2005) (Katoonizadeh et al., 2006), we investigated whether VEGFA could also promote generation of de novo hepatocytes from the expanded pool of BECs. If effective, VEGFA treatment could serve for the nearly 30% of severe cases of APAP overdose requiring liver transplantation (Yoon et al., 2016) as an alternative treatment to the drug NAC currently used in the clinic to neutralize APAP toxic metabolite NAPQI. Evidence of emergence of intermediate hepatocytes from BECs in severe intoxication in humans has been reported (Katoonizadeh *et al*., 2006), yet the process of BEC-to-hepatocyte conversion must be accelerated to become a viable and effective treatment for acute liver injuries, as proposed here with VEGFA mRNA-LNP. To better reflect clinical cases of human severe acute APAP intoxication in which hepatocyte senescence is consistently observed (Bird et al., 2018; Katoonizadeh *et al*., 2006), *Krt19*-Cre^ERT^, *R26*^LSL^ tdTomato mice were injected with AAV8-*Tbg*-p21 one week prior to APAP administration (Figure 3A). p21 was expressed in nearly 70% of hepatocytes (Figure S2A). APAP/p21 injury induced a ductular response strictly localized around the portal vein areas (Figure 3B), which was milder in the absence of AAV8-*Tbg*-p21 as expected. Two consecutive injections of VEGFA mRNA-LNP after APAP administration triggered remarkable BEC-to-hepatocyte conversion (Figure 3C). Numerous large clusters of tdTomato+ hepatocytes were evenly scattered in all liver lobes of VEGFA mRNA-LNP-treated mice, with a % of tdTomato+ areas 20-fold greater than in control LNP-treated mice (Figure 3C). Quantification of tdTomato+ hepatocytes was evaluated using flow cytometry analysis (Figure 3D) and was further adjusted for lineage tracing efficiency discrepancy (Figure 3D, Figure S2B). Adjusted % tdTomato+ hepatocytes were 5.6-fold greater in VEGFA mRNA-LNP-treated group compared to control LNP-treated mice (Figure 3D). BEC-derived hepatocytes were identified by co-expression of HNF4α (Figure 3E), and their BEC origin was supported by their proximity to bile ducts (Figure 3E). Interestingly, a few bi-phenotypic tdTomato+ BECs within a bile duct (Figure 3E, yellow arrowheads) were visualized by co-expression of the hepatocytic marker HNF4α as previously described (Deng *et al*., 2018). Periodic acid-Schiff staining of serial liver sections illustrated the ability of tdTomato+ hepatocytes to store glycogen as efficiently as adjacent tdTomato-hepatocytes (Figure 3F). Similar to the chronic CDE/p21 injury model, BEC-to-hepatocyte conversion did not occur in APAP injured mice in the absence of p21-induced hepatic senescence in control LNP-treated mice, while small clusters of tdTomato+ hepatocytes were consistently detected in all VEGFA mRNA-LNP-treated mice (Figure S2C, S2D). Our data demonstrate the ability of VEGFA mRNA-LNP treatment to trigger BEC-to-hepatocyte conversion that can be further amplified in the presence of hepatocyte senescence in an acute liver injury, suggesting VEGFA mRNA-LNP as an alternative treatment for severe APAP intoxication that would prevent liver failure and, thus, the need for transplantation.

**Figure 3.**
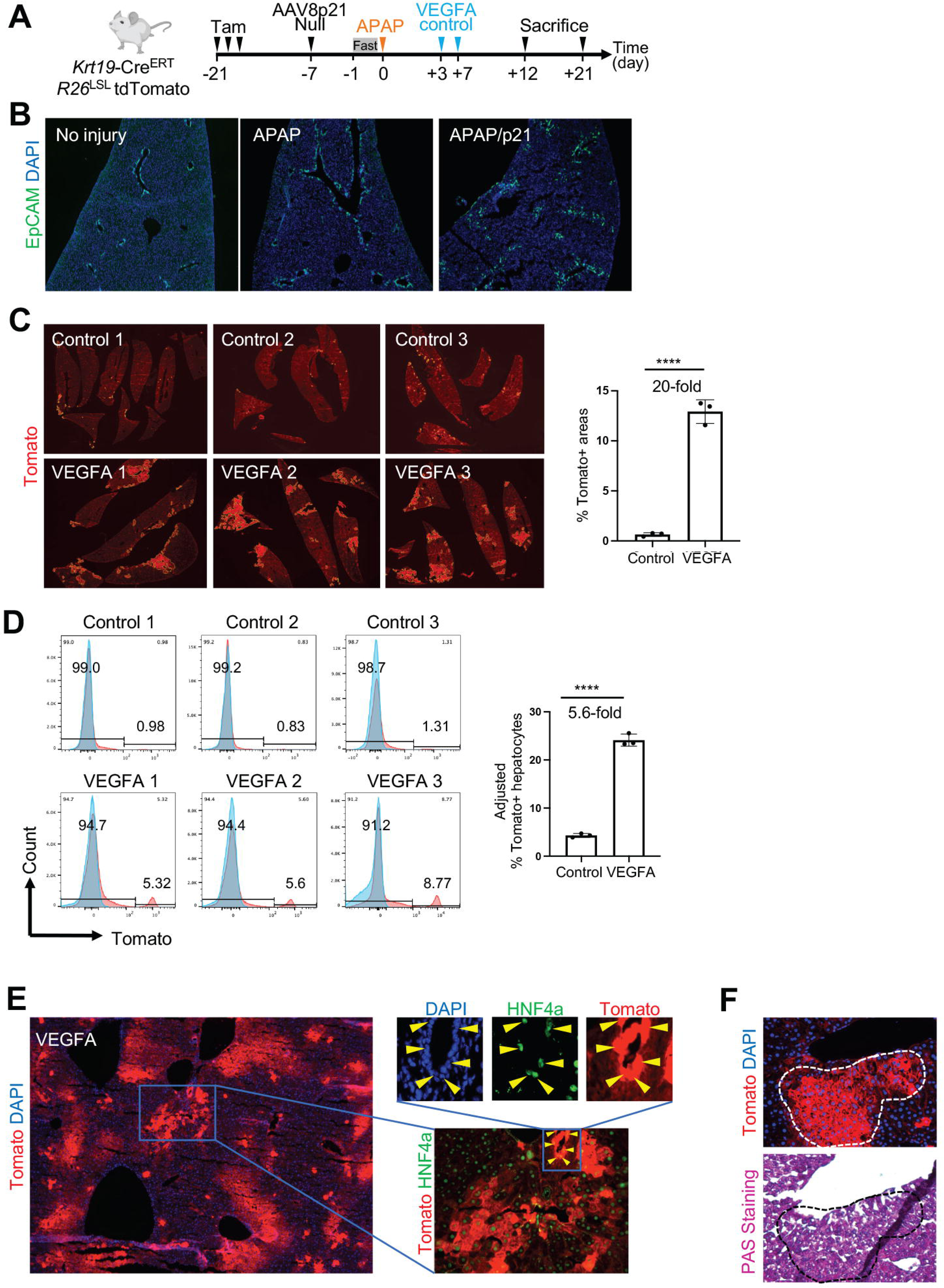
VEGFA mRNA-LNP administration induces BEC-to-hepatocyte conversion in APAP/p21-induced acute liver injury in mice. **(A)** Experimental design using the *Krt19*-Cre^ERT^, *R26*^LSL^tdTomato mice. AAV8-Tbg-p21 vector was administered to induce hepatocyte senescence. Injury was induced by a single intra-peritoneal injection of acetaminophen (APAP), followed by VEGFA mRNA-LNP or control Poly(C) RNA-LNP injections, 10 μg/20g body weight. **(B)** Representative immunofluorescent images showing comparison of EpCAM+ BECs observed in mice treated with APAP or APAP/p21. **(C)** Immunofluorescence microscopy images of tdTomato+ areas (outlined in yellow) in livers of mice given two injections of either control Poly(C) RNA-LNP (n=3) or VEGFA mRNA-LNP (n=3). The graph shows quantification of tdTomato+ area in both groups. **(D)** Histograms from flow cytometry of hepatocyte fraction isolated from mouse livers. Values of red histograms represent % tdTomato+ population in the hepatocyte fraction. Blue histograms represent hepatocytes from a control non-tdTomato background run simultaneously with experimental mouse. Bar graph shows the total % tdTomato+ hepatocytes calculated by extrapolating the lineage tracing efficiency as 100% across all mice. **(E)** Representative immunofluorescence microscopy images showing hepatocyte identity of tdTomato+ cells (arrows) with HNF4α staining (green) in the VEGFA mRNA-LNP-treated group. Close-up images show HNF4α+ cells within a tdTomato+ biliary duct. **(F)** tdTomato and PAS staining on serial sections of liver tissue demonstrating the ability of tdTomato+ hepatocytes to store glycogen. Data are presented as mean ± s.d. P values were determined by two-tailed Student’s t-test, *p< 0.05, **p< 0.01, ***p < 0.001, ****p<0.0001.

### VEGFA induces hepatocyte generation from KDR expressing cells in murine acute and chronic liver injury models and improves liver function

To further understand the regenerative role of VEGFA in mice, we identified the liver cell types expressing VEGFR2, also known as KDR, the main functional receptor for VEGFA (Holmes et al., 2007). As expected, endothelial cells, identified with CD31 expression, are the main liver cells expressing KDR as assessed by immunostaining on liver sections of non-injured mice (Figure S3A). Given that BECs have been reported to express KDR in rodent and human diseased livers (Fabris *et al*., 2006; Gaudio *et al*., 2006), we searched for KDR expression in BECs in the chronic CDE/p21 and acute APAP/p21 liver injury mouse models. In both models, we consistently observed expression of KDR in a subpopulation of BECs (Figure 4A, 4B) that was absent in control uninjured mice (not shown), suggesting that KDR marks a subset of BECs that may represent a facultative LPC that could be directly stimulated by VEGFA to generate hepatocytes. To test this hypothesis, we contracted the Jackson Laboratory to establish an inducible KDR lineage tracing line, *Kdr*-2A-Cre^ERT2^-2A-eYFP (Figure S3B) allowing cell fate mapping of KDR-expressing cells with time as well as marking KDR-expressing cells with YFP. Given that the 2A-Cre^ERT2^-2A-eYFP cassette was introduced downstream of the last exon of *Kdr* gene, this approach did not affect KDR expression, and therefore homozygous Cre/Cre survive, as opposed to the lethal KDR knockout mice (Shalaby et al., 1995). YFP is an accurate tool to track KDR expression as shown with overlapping co-staining marking endothelial cells (Figure S3C). In the absence of tamoxifen, in oil-treated mice, some reporter activity was detected in 5.42 ± 1% and 4.68 ± 2% of endothelial cells in females and males, respectively (Figure S3D, S3E, S4A), while leakiness in hepatocytes was almost nonexistent with rare tdTomato+ cells detected in <1:25,000 hepatocytes scanned in all liver lobes from 6 mice (Figure S4B). We were therefore confident to use the *Kdr*-2A-Cre^ERT2^-2A-eYFP; *R26*-tdTomato mice to map the hepatocyte fate of KDR expressing cells. Given the transient expression of KDR on BECs in response to injury, additional tamoxifen injections were included during the period of injury. In the CDE/p21 model, tamoxifen was injected 3 times over 5 days, starting from the fifth day of the diet (Figure 4C). The endothelial cell lineage mapping was robust in all mice, as virtually all endothelial cells were tdTomato+ (Figure S5A). We confirmed expression of KDR in a subset of BECs with the *Kdr* lineage tracing model by the presence of tdTomato+ BEC within biliary ducts (Figure 4D). Analyses of all liver lobes of each mouse revealed an even pattern of tdTomato+ cells representing endothelial cells (Figure 4E). Strikingly, some areas were much brighter (Figure 4E, areas within yellow dotted line), and were identified as tdTomato+ hepatocytes with co-expression of HNF4α (Figure 4F) and CD26 (Figure 4G). The tdTomato+ hepatocyte clusters were significantly larger and more numerous in VEGFA mRNA-LNP-treated mice with a 2.97-fold greater surface area (Figure 4E). As CD26 is a bile canalicular enzyme that depicts functional polarization of hepatocytes, its expression supports functional maturation of tdTomato+ hepatocytes derived from KDR+ cells. As noticed for the *KRT19* lineage tracing model, extent of DR was similar between the two groups (Figure S5B), most likely due to greater BEC-to-hepatocyte conversion in the VEGFA mRNA-LNP-treated group. Importantly, we confirmed that VEGFA mRNA-LNP treatment significantly reversed steatosis as compared to control-LNP treated mice (Figure 4H) as seen in the CDE/p21-treated *Krt19* lineage tracing mice.

**Figure 4.**
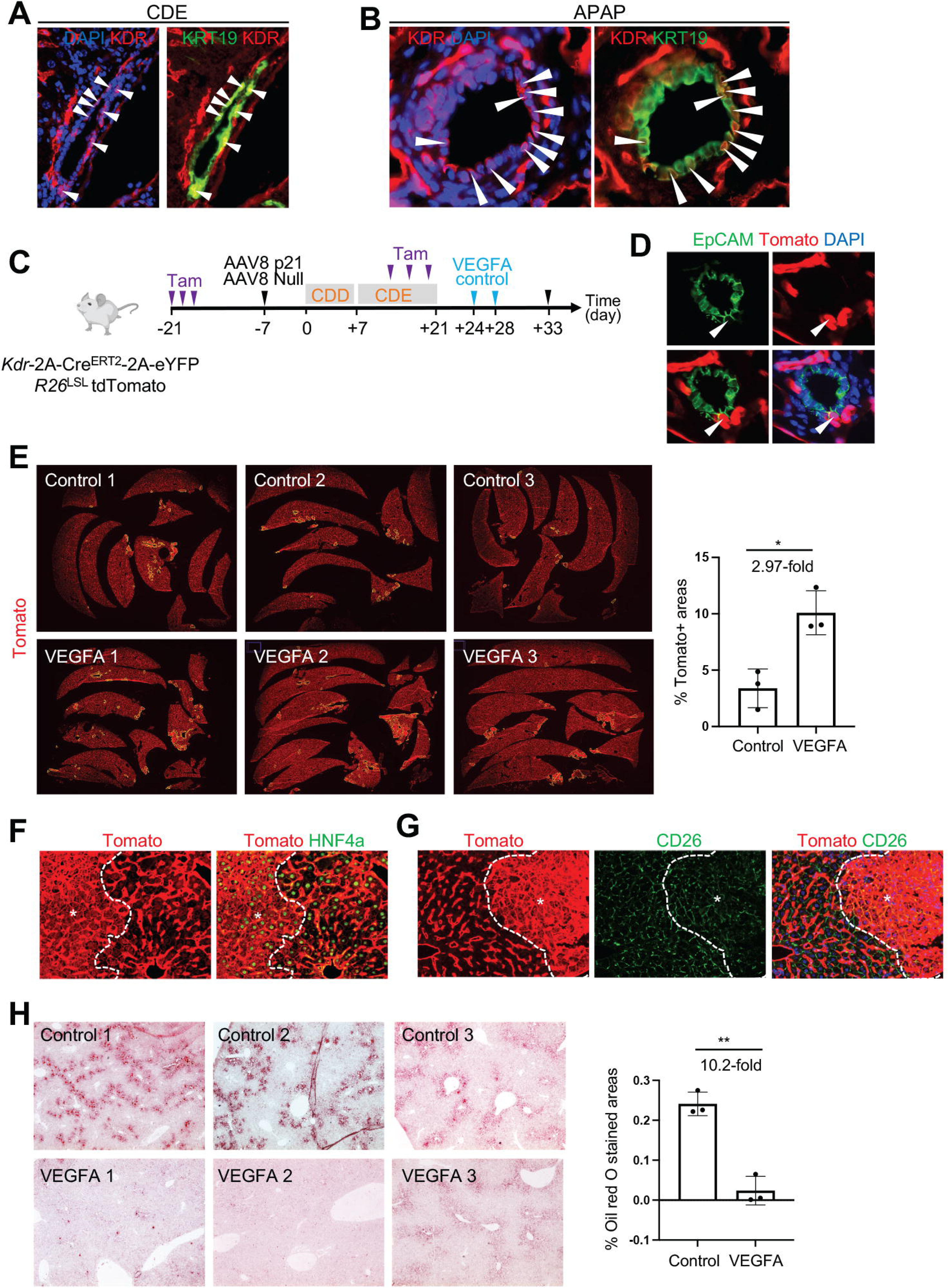
VEGFA mRNA-LNP administration induces BEC-to-hepatocyte conversion in CDE/p21-induced chronic liver injury in *Kdr*-2A-Cre^ERT2^-2A-eYFP mice. **(A, B)** Expression of KDR in BECs from the CDE- and APAP-induced liver injuries in the presence of AAV8-Tbg-p21. KDR expression (red) was observed in some of the BECs (green) depicted by arrowheads. **(C)** Scheme showing key interventions in the experimental design using the *Kdr*-2A-Cre^ERT2^-2A-eYFP, *R26*^LSL^tdTomato mice to track any cell that expresses KDR using CDE/p21 injury model. Additionally, three injections of tamoxifen were given on alternate days in the second week of injury to trace any BECs expressing KDR. **(D)** Representative close-up images of a bile duct showing tdTomato+ EPCAM+ BECs. (E) Immunofluorescence microscopy images showing tdTomato+ areas (outlined yellow) in livers of mice given 2 injections of either control Poly(C) RNA-LNP (n=3) or VEGFA mRNA-LNP (n=3). The bar graph shows the quantification of the tdTomato+ area in the two groups. **(F, G)** Representative immunofluorescence images depicting hepatocyte identity of tdTomato+ cells (outlined and identified with *) with HNF4α staining (green), a hepatocyte marker, or CD26 staining (green), a mature hepatocyte marker expressed on the canalicular face of the hepatocytes. **(H)** Representative brightfield images of Oil Red O-stained liver tissues showing lipid accumulation in the liver. The bar graph shows the quantification of the % Oil Red O-stained area averaged from at least three different fields in each mouse in the two groups (n=3 per group). Numerical data are presented as mean ± s.d. P values were determined by two-tailed Student’s t-test, *p< 0.05, **p< 0.01.

The findings were further supported in the acute APAP/p21-induced liver injury (Figure 5). Tamoxifen was additionally injected 3 times, 1 day prior and 1 and 3 days after APAP administration to capture the emerging KDR+ BECs upon injury (Figure 5A). VEGFA mRNA-LNPs significantly promoted emergence of bright tdTomato+ areas (Figure 5B, 6.7-fold increase) identified as hepatocytes with co-staining for HNF4α (Figure 5C) and CD26 (Figure 5D) as compared to control-LNP-treated mice. Quantification of the KRT7+ BEC areas in both control-LNP- and VEGFA mRNA-LNP-treated mice confirmed again that these numbers were not significantly different (Figure S5C) as found in the CDE/p21 model. These data are reminiscent to observations made in humans following acetaminophen intoxications in which the ductular reaction decreases as BEC-to-hepatocyte conversion occurs (Katoonizadeh *et al*., 2006). To capture early events of KDR+ BEC conversion into hepatocytes during APAP/p21 injury, we examined injured *Kdr*-2A-Cre^ERT2^-2A-eYFP mice (in absence of lineage tracing, Figure S5D) three days following APAP administration (Figure 5D). We detected many YFP+ BECs in areas indicative of early regeneration featured by endothelial cell ablation and newly generated hepatocytes with intact DAPI staining (Figure 5E, area delineated with dotted line). The presence of large YFP+ hepatocyte-like cells (Figure 5E, arrows) suggests their differentiation from YFP+KDR+ BECs (Figure 5E, arrowheads). Observation of YFP+ hepatocyte-like cells was most likely possible because of the longer half-life of YFP protein compared to that from KDR protein.

**Figure 5.**
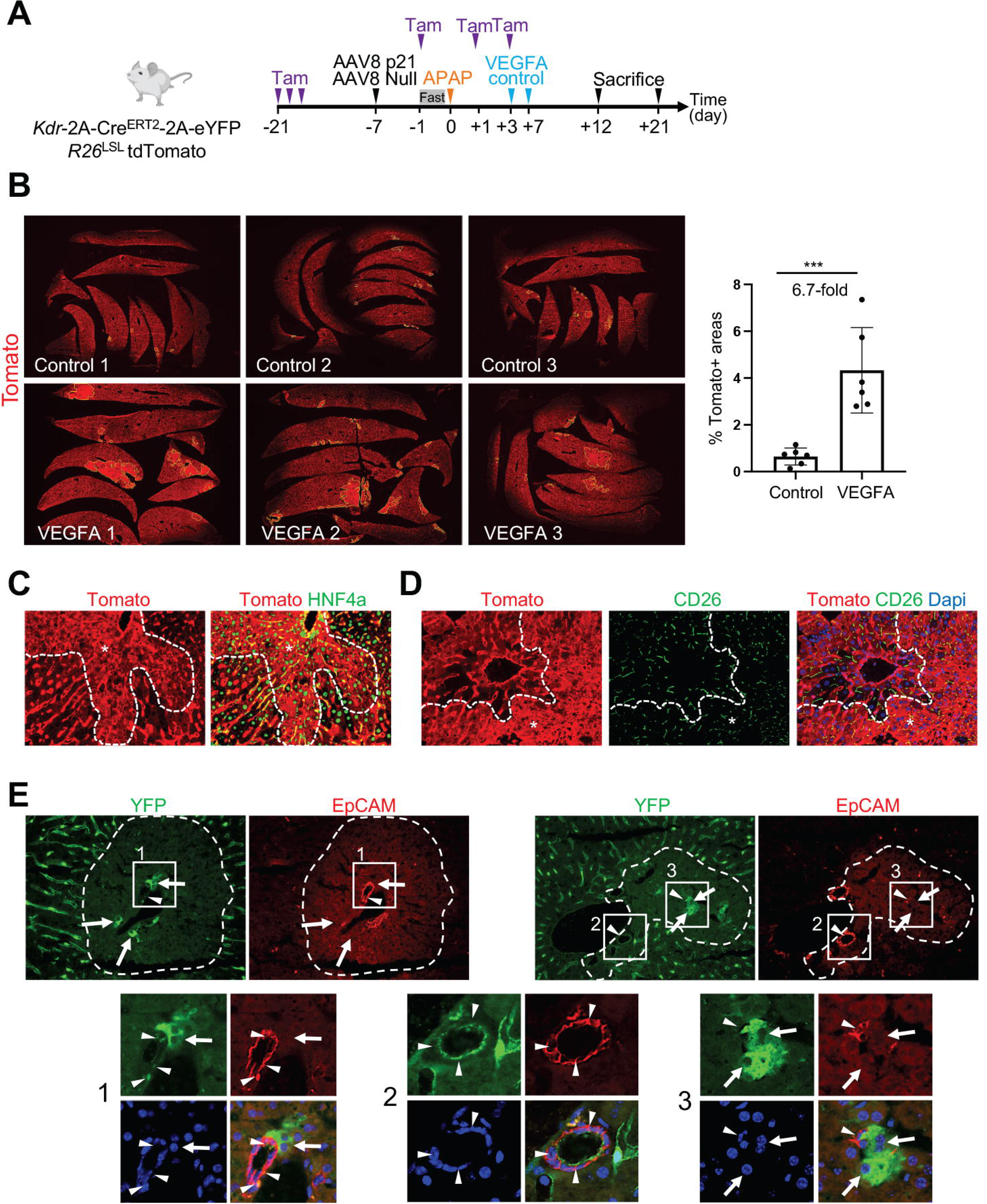
VEGFA mRNA-LNP administration induces BEC-to-hepatocyte conversion in APAP/p21-induced acute liver injury in *Kdr*-2A-Cre^ERT2^-2A-eYFP mice. **(A)** Scheme showing key interventions in the experimental design using the *Kdr*-2A-Cre^ERT2^-2A-eYFP, *R26*^LSL^tdTomato mice to track any cell that expresses KDR in APAP/p21 injury model. Three injections of tamoxifen are given on alternate days around injury to trace any BEC expressing KDR in response to APAP injury. **(B)** Immunofluorescent microscopy images showing tdTomato+ areas (outlined yellow) in livers of control firefly luciferase mRNA-LNP- (n=6) or VEGFA mRNA-LNP- (n=6) treated mice. The bar graph shows quantification of tdTomato+ area in the two groups. **(C, D)** Representative immunofluorescent images depicting hepatocyte identity of tdTomato+ cells (outlined and identified with *) with HNF4α staining (green), a hepatocyte marker or CD26 staining (green), a mature hepatocyte marker that is expressed on the canalicular face of the hepatocytes. **(E)** Immunofluorescent images showing expression of YFP in liver cells of *Kdr*-2A-Cre^ERT2^-2A-eYFP mice three days after acute liver toxicity by intraperitoneal APAP injection (400 mg/kg to male or 500 mg/kg to female). The fields of interest have been enlarged below. Arrowheads depict BECs co-expressing YFP (green) and EpCAM (red) while arrows depict large intermediate hepatocyte-like cells expressing YFP (green). Numerical data in the figure are presented as mean ± s.d. P values were determined by two-tailed Student’s t-test for comparison between two groups, ***p < 0.001.

Overall, findings from complementary *Krt19*- and *Kdr*-lineage tracing models demonstrate the conversion of KDR+ expressing cells, most likely BECs, into hepatocytes which is significantly augmented after VEGFA mRNA-LNP administration, a therapeutic strategy that could be leveraged in the clinic to treat acute and chronic liver diseases.

### Identification of KDR-expressing BECs with evidence of BEC-to-hepatocyte conversion from explanted cirrhotic liver specimens with end-stage liver disease (ESLD) and Child-Pugh score B/C

To investigate the clinical relevance of our findings from the zebrafish and mouse models, we sought to identify KDR-expressing BECs associated with evidence of BEC-to-hepatocyte conversion in specimens from human cirrhotic livers with ESLD recovered from patients with NASH cirrhosis or alcoholic cirrhosis and terminal liver failure (n = 5) (Child-Pugh B and C), as well as three normal specimens (Table S1). Histopathological examination confirmed that all five diseased specimens exhibited features of ESLD such as hepatocyte-containing regenerative nodules surrounded with fibrotic septa (Figure S6A, S6B) associated with various degrees of steatosis (Figure S6C) and ductular reaction (Figure S6D). Numerous KRT7+ cells were also found near the margin of, as well as within, the regenerative nodules (Figure 6A), suggesting the presence of KRT7+ intermediate hepatocyte-like cells derived from KRT7+ BECs as previously speculated in humans (Sancho-Bru et al., 2012). As expected, expression of glutamine synthetase (GS) in hepatocytes around central vein areas was detected in diseased specimens as found in normal specimens (Figure S7A,B,D). However, aberrant GS expression was also found in KRT7+ BECs within the DR and in the adjacent transitioning KRT7+ hepatocytes as well as their neighbors and most likely progeny KRT7-hepatocytes (Figure 6B) as described (Fleming and Wanless, 2013; Stueck and Wanless, 2015). GS expression became weaker as hepatocytes were further away from the DR, toward the center of the regenerative nodule (Figure 6B, asterisk). Large GS+ hepatocyte-like cells were often seen budding from strings of smaller KRT7+ BECs (Figure 6B, arrows), an observation that is reminiscent of the hepatocytic buds described two decades ago by Stueck and Wanless (Stueck and Wanless, 2015; Wanless *et al*., 2000). Importantly, while KDR expression was not detected in BECs from normal specimens (Figure S7C), numerous KRT7+ KDR+ BECs were identified histologically in 2 out of 5 cirrhotic specimens (specimens HH125 and HH121, Figure 6C, frames 4 and 5, arrowheads). Strikingly, KRT7- KDR+ hepatocyte-like cells (Figure 6C, frames 2 and 3, arrows) were seen budding from the strings of KRT7+KDR+ BECs (Figure 6C, arrowheads), as observed in mice in the APAP/p21 injury model (*Kdr*-2A-Cre^ERT2^-2A-eYFP) in which YFP+ hepatocytes were found adjacent to the YFP+KDR+ BECs (Figure 5E, arrows). Expression of cytoplasmic KDR was also found in a subset of hepatocytes adjacent to KDR+ BECs (Figure 6C, frame 1, arrows). We further validated KDR expression in purified human hepatocytes from 9 explanted cirrhotic livers (NASH or alcoholic) patients that underwent liver transplantation due to decompensated hepatic function (Child-Pugh score B/C) and compared to normal human liver specimens (Table S1) as previously described (Gramignoli et al., 2012) at the transcript (Figure 6D, 9 specimens) and protein levels (Figure 6E, 6 specimens). We first validated the hepatocyte identity of our purified cell preparation with the similar levels of mRNA HNF4α transcript representing promoter P1- and P2-derived HNF4α isoforms in the 3 types of specimens including hepatocytes purified from normal and both Child-Pugh score B or C diseased specimens (Figure 6D). As expected from our previous studies, protein levels of P1-derived adult HNF4α isoforms were critically downregulated in diseased hepatocytes compared to those in normal hepatocytes (Heps) or normal liver tissue (Aguila et al., 1997; Guzman-Lepe et al., 2018; Nishikawa et al., 2015). Importantly, KDR and KRT7 transcript as well as protein levels were upregulated in purified diseased hepatocytes (Fig 6d, 6e, Extended data Figure 7E) compared to normal human hepatocytes. Of note, diseased hepatocytes expressed the distinct 150 KDa non-glycosylated isoform of KDR (Fig 6e), that was not detected in purified normal hepatocytes nor the normal liver tissue specimens. These results support the specificity of KDR to diseased hepatocytes as opposed to potential contaminant endothelial cells from the preparation of purified hepatocytes. Indeed, the 230 KDa isoform of KDR was the only variant detected in normal liver tissues that include endothelial cells. The 150 KDa KDR isoform is known to remain in the cytoplasm as an inactive receptor (Takahashi and Shibuya, 1997) which is consistent with the cytoplasmic staining of KDR observed in specimen sections (Figure 6C, arrowheads). Although, we cannot conclude about the function of non-glycosylated isoform of KDR in hepatocytes, it serves here as a surrogate lineage tracing mapping the hepatocytic cell fate of KDR-expressing BECs. In line with the notion of VEGFA as a therapeutic target for advanced liver diseases, a recent study has shown a significant correlation between high levels of serum VEGF and lower fibrosis score in a NAFLD cohort (Papageorgiou et al., 2017), suggesting a protective effect of VEGFA against progression of the disease. Yet, the study indicates that levels of serum VEGF tend to decrease when NASH develops, offering a potential therapeutic intervention for VEGFA mRNA-LNP to mitigate the liver disease.

**Figure 6.**
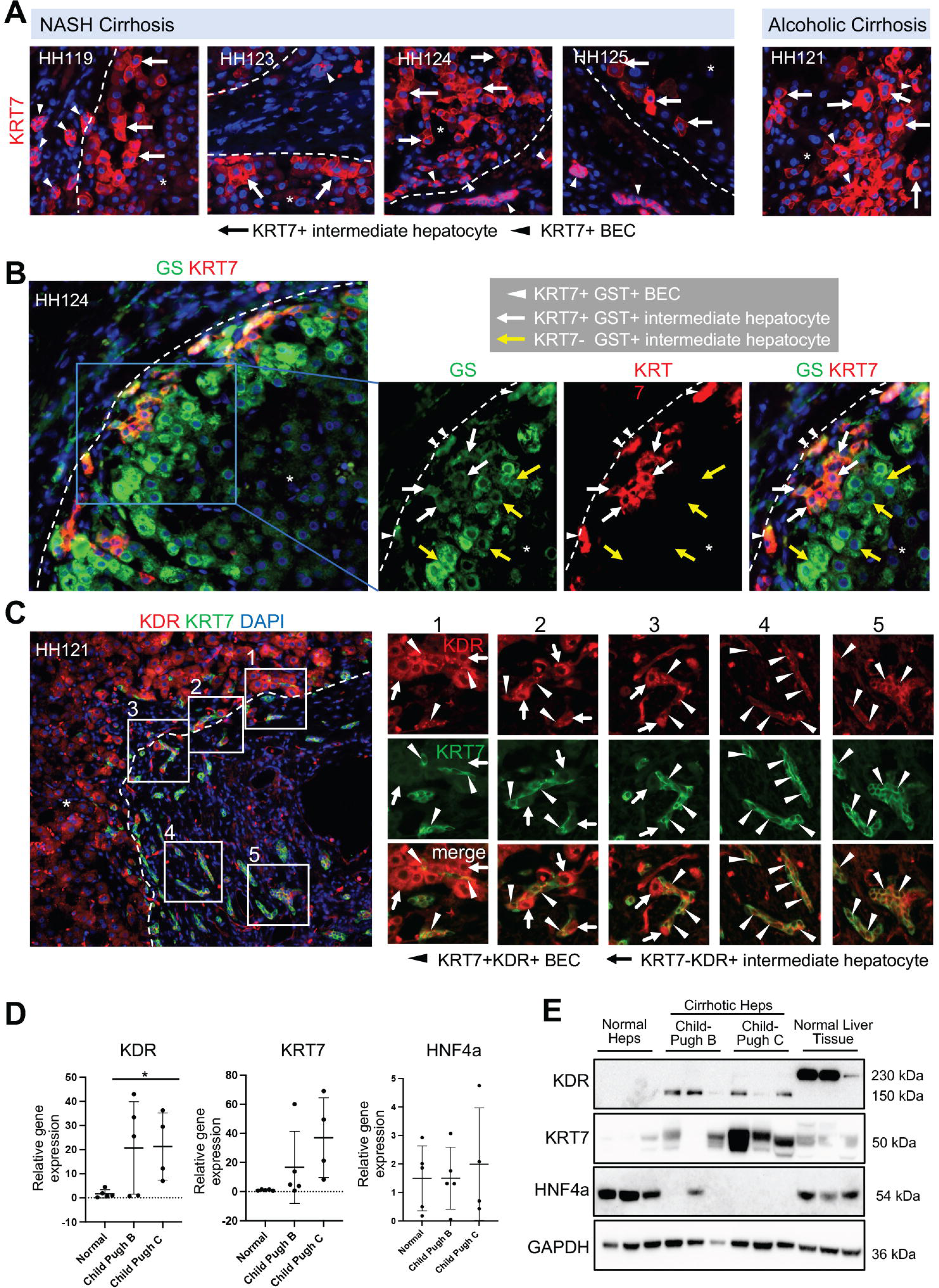
Evidence of BEC-to-hepatocyte conversion and KDR expression in human liver samples from non-alcoholic steatohepatitis (NASH) cirrhosis and alcoholic cirrhosis with ESLD patients. **(A)** Immunofluorescent images showing small cuboidal KRT7+ BECs (arrowheads) and KRT7+ intermediate hepatocyte-like cells (arrows) largely present in the regenerative nodules (*) close to the fibrous septa (outlined). **(B)** Immunofluorescence images showing KRT7 (red) and GS (green) staining in human cirrhotic liver specimens. The white arrowheads depict GS+ KRT7+ BECs (yellow cells) possessing a duct cell-like morphology. The white arrows depict the rare GS+ KRT7+ intermediate hepatocyte-like cells, while yellow arrows depict GS+ KRT7- intermediate hepatocyte-like cells. **(C)** Immunofluorescent images of cirrhotic liver specimen showing fibrous septa stained for KRT7+ BECs (green) and KDR+ sinusoidal endothelial cells (red). Numerous KRT7+ KDR+ (arrowheads) BECs can be observed (magnified images) along with larger KDR+ intermediate hepatocyte-like cells (arrows) surrounding the KRT7+ KDR+ BECs. **(D)** Relative gene expression of KDR, KRT7, HNF4α determined in hepatocytes (Heps) isolated from normal (n=5) or cirrhotic human livers, Child-Pugh B (n=5) and Child-Pugh C (n=4). **(E)** Western blots showing protein expression of KDR, KRT7, HNF4α in hepatocytes isolated from normal or cirrhotic human livers Child-Pugh B (n=3) and Child-Pugh C (n=3) analyzed above in panel d. GAPDH serves as endogenous control. Numerical data in the figure are presented as mean ± s.d. P values were determined by two-tailed Student’s t-test for comparison between two groups, *p < 0.05.

Altogether, identification in human ESLD specimens of KDR+ BECs associated with evidence of BEC-to-hepatocyte conversion supported by the presence of KRT7+GS+ hepatocytes and KDR+ hepatocyte-like cells budding from KRT7+GS+ BECs and KRT7+KDR+ BECs, respectively, provides a potential clinical application for VEGFA mRNA-LNP to stimulate BEC-driven liver repair for human liver disease intervention.

## Discussion

Hepatocyte-driven repair fails in the face of severe acute injury or years of build-up of chronic liver damage, unveiling the possible clinical benefit of an alternative repair mechanism mediated by BECs. Decades of literature from analyses of human chronic liver disease specimens identified hepatocytes within regenerative nodules or adjacent to portal vein triads, that harbor BEC marker expression such as KRT7 or EpCAM (Katoonizadeh *et al*., 2006; Limaye et al., 2008; Vandersteenhoven et al., 1990; Yoon *et al*., 2011) as well as BECs within the DR turning on a hepatocyte signature with concomitant expression of HNF4α, HNF6 or GS (Falkowski *et al*., 2003; Fleming and Wanless, 2013; Haque *et al*., 1996; Hytiroglou and Theise, 2018; Limaye *et al*., 2008; Stueck and Wanless, 2015; Wanless *et al*., 2000). Given the clinical context of these human advanced chronic liver disease specimens in which proliferation of senescence gene-expressing hepatocytes is exhausted (Bird *et al*., 2018; Katoonizadeh *et al*., 2006; Marshall *et al*., 2005; Richardson *et al*., 2007), these observations suggest the BEC origin of the bi-phenotypic hepatocyte-like cells. More recent lineage tracing studies in zebrafish and mouse models of liver injuries have recapitulated the generation of DR and demonstrated BEC-to-hepatocyte conversion when hepatocyte proliferation is compromised (Choi *et al*., 2014; Deng *et al*., 2018; He *et al*., 2014; Lu *et al*., 2015; Manco *et al*., 2019; Raven *et al*., 2017; Russell *et al*., 2018). However, the presence of DR in human advanced chronic liver diseases has been associated with poor prognosis (Sancho-Bru *et al*., 2012), casting a doubt on the BEC ability to promote liver repair and thus questioning their clinical benefit as facultative progenitor cells. Indeed, BECs within DR release profibrogenic factors that may instead aggravate the chronic liver disease features (Glaser et al., 2009; Kaur et al., 2015). Yet, a recent study showed that aberrant GS positivity in hepatocytes adjacent to portal tracts is significantly associated with regressed cirrhosis in humans (Hadi *et al*., 2020), suggesting, in contrast, a positive clinical outcome from BEC-derived hepatocytes in resolving human cirrhosis. One of the most convincing studies in humans illustrating the ability of BECs to convert into hepatocytes was attempted by lineage tracing the BECs among the DRs in human cirrhotic livers using mutational analysis in mitochondrial DNA encoding cytochrome c oxidase enzyme, and showed the descent of hepatocytes within monoclonal regenerative nodules from adjacent BEC-associated DRs (Lin *et al*., 2010). Altogether, experimental animal model studies combined with analyses of human specimens raise the possibility of leveraging the naturally occurring DR and associated BEC-to-hepatocyte conversion as a therapy if the BEC-driven repair mechanism could be harnessed. This would have a tremendous impact on the treatment of acute and chronic liver diseases and would circumvent liver transplantation and accompanying challenges and complications.

The current challenge for the efficient therapeutic use of BECs is the inability to reliably identify a true progenitor population among them and to, thus, define a druggable pathway that would accelerate their differentiation into functional hepatocytes. Here, using complementary mouse and zebrafish liver injury models, we demonstrate that VEGFA potentializes BEC-driven liver regeneration by promoting BEC-to-hepatocyte conversion with a 5-fold increase on average. The presence of KDR-expressing BECs in advanced cirrhotic human liver specimens as well as adjacent KDR-expressing hepatocytes, most likely progeny of the KDR-expressing BECs, suggests that BEC-to-hepatocyte conversion occurs in humans and is possibly mediated through KDR activation on KDR-expressing BECs. Therefore, if augmented with VEGFA mRNA-LNP, BEC-driven liver repair may become an efficient therapy in humans to prevent progression of the chronic liver disease and promote its regression, or to prevent liver failure in acute disease overcoming the necessity for liver transplantation.

The precise identity of BEC liver progenitor cells remains elusive. Although, there is no evidence for a direct relationship between fetal progenitor hepatoblasts and adult liver progenitor cells, they do share the same clonogenic features with capacity to differentiate into hepatocytes and BECs, as well as share several cell surface markers including DLK1, CD133 and EpCAM (Dorrell et al., 2011; Jensen et al., 2004; Kamiya et al., 2009; Okabe et al., 2009; Qiu et al., 2011; Rountree et al., 2007; Schmelzer et al., 2007; Suzuki et al., 2008; Tanaka et al., 2009; Yovchev et al., 2008) (Ko et al., 2020; So et al., 2020). However, since these markers are expressed on all BECs in adult livers, it is therefore difficult to identify the clonogenic progenitors amongst the non-clonogenic BECs. Other markers such Lgr5 (Huch et al., 2013), TROP2 (Okabe *et al*., 2009), Foxl1 (Shin et al., 2011), and ST14 (Li et al., 2017) have been reported to define BEC progenitors, yet their cell fate mapping during regeneration have not been fully performed or the ability of their progeny to restore liver function fully demonstrated. We previously identified KDR as a distinct cell surface marker expressed by a fetal liver progenitor (Goldman et al., 2013) that is conserved in both murine and human fetal liver. Our studies using the mouse and the human embryonic stem cell (ESC) differentiation system revealed that KDR+ fetal hepatic progenitors are *bona fide* hepatoblast precursors and that KDR activation is required for hepatic specification of the human ESC-derived KDR+ hepatic progenitor. This prompted us to investigate the possibility of re-expression of KDR on a subset of adult BECs upon injury and its activation with VEGFA as a therapeutic means to reverse the liver disease. In accordance with the few studies reporting KDR expression on BECs (Dianat *et al*., 2014; Fabris *et al*., 2006; Gaudio *et al*., 2006), we identified KDR+ BECs in mouse liver injury models as well as the ability of KDR+ cells to generate hepatocytes using the *Kdr* lineage tracing model (see graphical abstract). Importantly, as found in mice, KDR expression was also detected on BECs in human cirrhotic ESLD specimens as well as on hepatocytes budding from strings of KRT7+ BECs. Presence of KDR+ hepatocytes in human cirrhotic ESLD specimens or YFP+ hepatocytes in injured livers in mice carrying the YFP reporter under the *Kdr* promoter supported their identity as progeny of KDR+ BECs. This study introduces KDR+ BEC as a novel facultative liver progenitor cell. Moreover, the use of clinically relevant, non-integrative mRNA-LNP, whose safety has been clinically validated (Weissman, 2015) and further supported by the current COVID-19 vaccines, to transiently deliver VEGFA in the liver may potentially have key clinical significance to treat chronic and acute liver diseases.

## STAR★Methods

### Key Resources Table

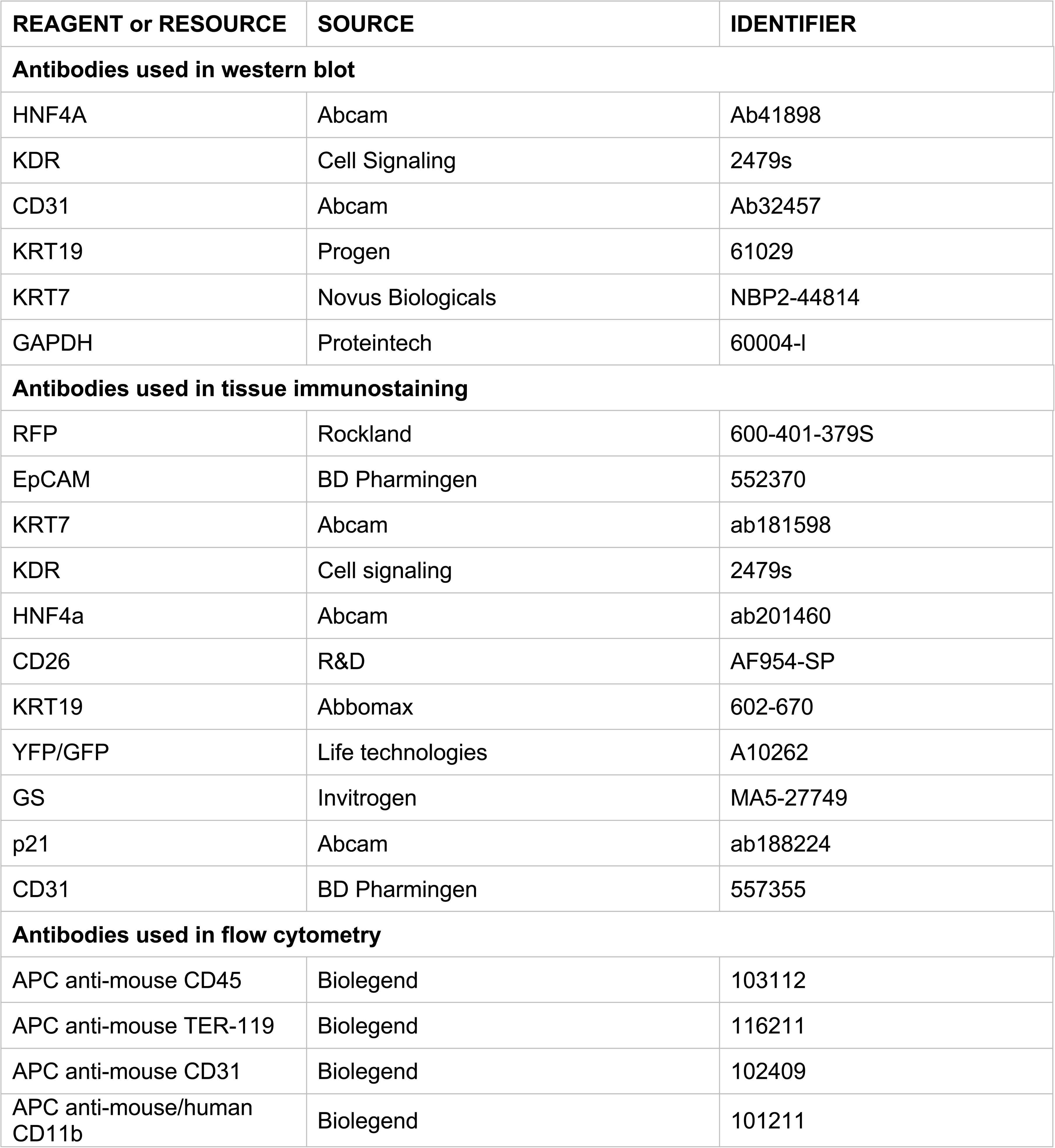
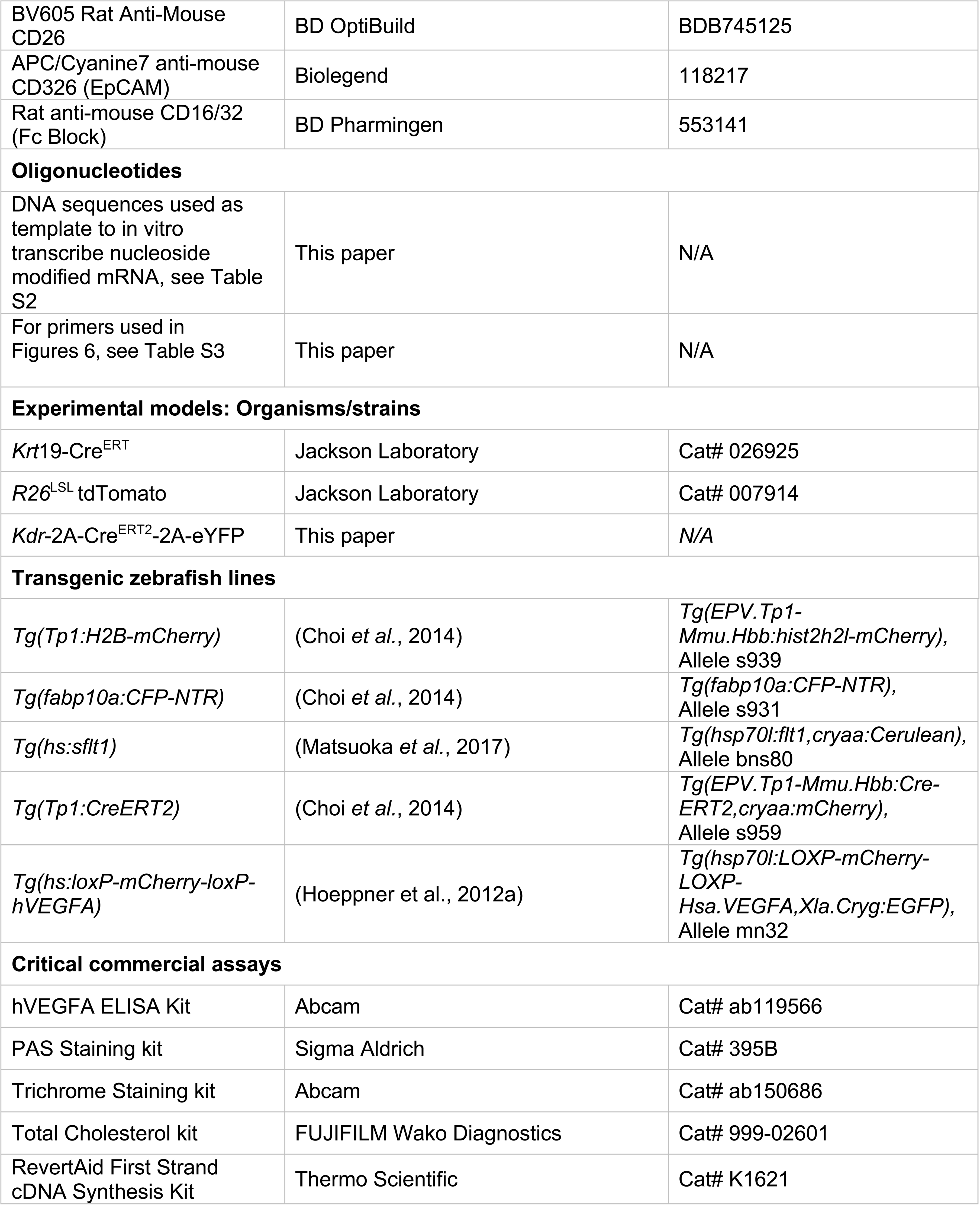
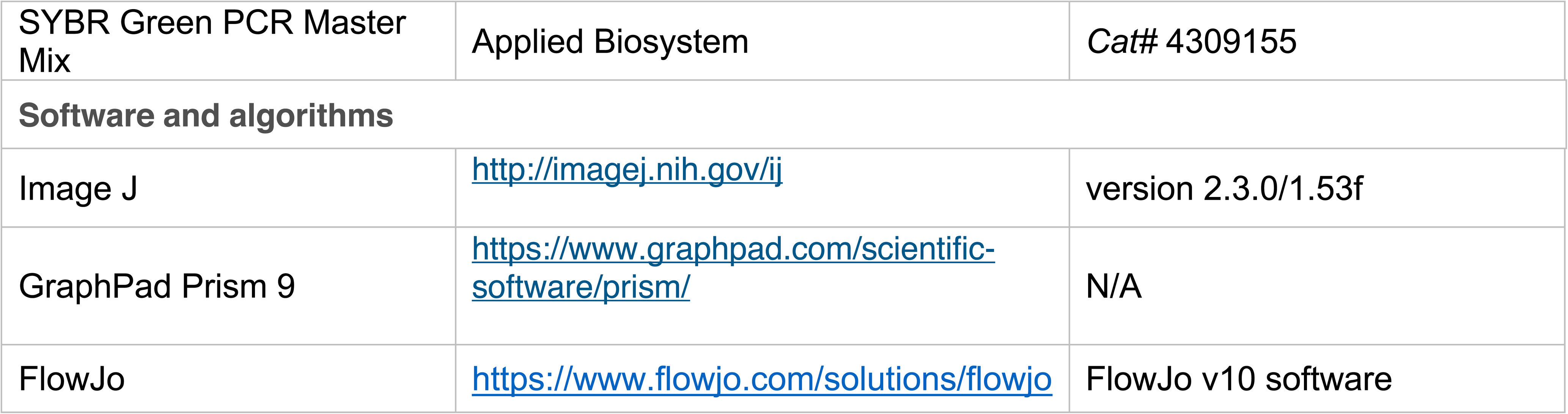

### Resource availability

#### Lead contact

Further information and requests for resources and reagents should be directed to and will be fulfilled by the lead contact, Valerie Gouon-Evans.

### Materials availability

This study did not generate new unique reagents.

### Experimental model and subject details

#### In vivo studies

All animal studies were approved by the Boston University IACUC and were consistent with local, state, and federal regulations as applicable. All experiments were done in an age and sex-controlled fashion unless otherwise noted. *Krt*19-Cre^ERT^ (Means et al., 2008) and *R26*^LSL^ tdTomato lines were obtained from Jackson Labs. Animals were housed under standard conditions with a 12-hour day/night cycle, in a pathogen-free environment with access to food and water ad libitum.

#### Generation of *Kdr*-2A-Cre^ERT2^-2A-eYFP lines

The Kdr knock-in mouse allele was generated using direct delivery of CRISPR-Cas9 reagents to mouse zygotes. Nucleotide changes in the mouse Kdr gene (Ensembl Gene UID ENSMUSG00000062960) were introduced in the Ensembl Kdr-201 transcript (ENSMUST00000113516.1). The mutant allele inserts the *Kdr*-P2A-CRE.ERT2-P2A-YFP.

##### Gene Editing Design

Analysis of genomic DNA sequence surrounding the target region, using the Benchling (www.benchling.com) guide RNA (gRNA) design tool, identified one gRNA sequence with a suitable target endonuclease site in exon 30. Predicted off-target sites for the gRNA (sgRNA2-ACCTCCTGTTTAAATGGAAG) were identified using the specificity model developed and documented by Hsu and colleagues and embedded in the Benchling gRNA design tool(Hsu et al., 2013). Design considerations for off-target editing were as follows: higher off-target editing risk was considered when the “Score” is >2.0 with a canonical SpCas9 PAM (NGG), >50.0 with a chromosomally unlinked non-canonical PAM (NAG), or >20.0 with a linked non-canonical PAM yet in an annotated protein coding region. These “Score” thresholds were based on empirical targeted sequencing analysis of 20 different predicted off-target loci in 15 different mouse zygote gene editing experiments.

##### sgRNAs and plasmid donor DNA repair templates

*Streptococcus pyogenes* Cas9 (SpCas9) mRNA was purchased from Trilink (Product# L-7606, Trilink, USA). sgRNAs were synthesized as described(Qin et al., 2015). The sequence of the double stranded DNA donor plasmid, that functions as the DNA repair template, was synthesized by Genscript. Features included asymmetric homology arm lengths of 3kb and 2.2kb, flanking the 2.8kb insertion sequence.

##### In vitro cell line gRNA evaluation

Using a mouse kidney cell line and a T7 endonuclease cleavage assay, individual gRNA duplexes were evaluated for their ability to direct SpCas9 protein to the target locus and induce a double stranded DNA cleavage event. The immortalized mouse kidney cell line MK4 (a gift from Yueh-Chiang Hu, University of Cincinnati) was infected with a lentivirus construct expressing a SpCas9-P2A-EGFP transgene. Infected cells were selected for high GFP expression using flow cytometry. 200 ng of the sgRNA duplexes were transfected into 50,000 MK4-SpCas9-P2A-EGFP transgenic cells in a 24 well plate using MessengerMax (Product# LMRNA015, ThermoFisher-Invitrogen, USA) according to manufacturer’s instructions; 200 ng mCherry mRNA (Product# L-7203, Trilink Biotechnologies, USA) was transfected in a separate well as a control. 24-48 hrs post-transfection, DNA lysates were prepared, and PCR was performed with primers flanking the DNA cleavage target site. PCR products were evaluated on agarose gels.

For evaluating the gRNA activity, the presence of small insertions/deletions (INDELs) in the population of PCR products was detected by denaturation and re-annealing of the PCR product and subsequent treatment with T7 endonuclease (Product# E3321, New England Biolabs, USA). T7 endonuclease cleaves heteroduplex DNA resulting from various INDELs in the population. Unique DNA cleavage products resulting from T7 endonuclease activity were resolved by agarose gel electrophoresis and compared to control transfection PCR products.

##### Preparation of gene editing reagents for mouse embryo microinjection

The gene editing reagents were prepared for microinjection as described previously(Qin *et al*., 2015). Briefly, 500ng/ul of Cas9 mRNA, 500ng/ul of sgRNA and 500ng/ul donor plasmid were assembled in DNase/RNase free TE buffer, centrifuged and the supernatant collected and delivered for microinjection, all kept on ice or at 4°C.

##### Zygote isolation, microinjection and embryo transfer

All animal work was approved by the Jackson Laboratory Animal Care and Use Committee and adhered to the standards of the Guide for the Care and Use of Laboratory Animals set forth by the NIH. Fertilized mouse embryos were generated via natural mating and cultured as described previously(Qin *et al*., 2015). C57BL/6J (Stock# 000664, The Jackson Laboratory, USA) donor female mice (3–4 weeks of age) were superovulated by administration of 5 IU of pregnant mare serum gonadotrophin (PMSG) via intraperitoneal (ip) injection (Product# HOR-272 ProSpec, Israel) followed 47 hr later by 5 IU (ip) human chorionic gonadotrophin (hCG) (Product# HOR-250, ProSpec, Israel). Immediately post-administration of hCG, the female was mated 1:1 with a C57BL/6J stud male and 22 hr later checked for the presence of a copulation plug. Female mice displaying a copulation plug were sacrificed, the oviducts excised, and embryos collected. Microinjection was performed as described(Qin *et al*., 2015). In brief, zygotes were microinjected on a Zeiss AxioObserver.D1 using Eppendorf NK2 micromanipulators in conjunction with Narashige IM-5A injectors. Embryos were immediately transferred into B6Qsi5F1 pseudopregnant female mice, a F1 hybrid strain produced by breeding C57BL/6J female mice with the inbred Quackenbush Swiss line 5 (QSi5) mouse strain.

##### Sequence characterization of Founder and N1 mice

PCR primers flanking the gRNA target site, but outside the repair template homology arms, were used to amplify the region of interest from founder progeny. PCR amplicons were subjected to Sanger sequencing to identify founders with precise desired changes. Founders carrying desired mutations were bred with C57BL/6J mice (Stock #000664, The Jackson Laboratory, USA) to produce N1 progeny. N1 animals were confirmed by the same PCR-Sanger sequence analysis the founder population was subjected to. In contrast to founders, N1 animals are obligate heterozygotes, with one allele derived from an un-manipulated chromosome, enabling deconvolution of the two alleles in the mixed PCR amplicon.

#### Collection of human liver specimens and hepatocyte isolation

Adult human liver cells were obtained from the Human Synthetic Liver Biology Core at the Pittsburgh Liver Research Center. The Institutional Review Board at the University of Pittsburgh has approved our protocol and given the Not Human Research Determination. The IRB# STUDY20090069. Hepatocytes were isolated using a three-step collagenase digestion technique as previously described (Gramignoli *et al*., 2012). Cell viability was assessed after isolation as previously described using trypan blue exclusion, and only cell preparations with viability > 80% were used for the analysis. Information about the human liver specimens and hepatocytes used in this study can be found in Table S1.

### Method details

#### Acetaminophen (APAP) induced acute injury model

Mice were fasted overnight for a period of 14-15h prior to APAP injections. APAP was administered at a concentration of 20mg/ml, which was freshly dissolved in sterile PBS at 56°C. After complete dissolution, the stock was cooled down to room temperature. A single intraperitoneal injection was administered to male and female mice which received either 300mg/kg or 500mg/kg body weight of APAP, respectively, unless otherwise mentioned. Mice were maintained on normal chow diet and water ad libitum after APAP injections.

#### CDE diet induced chronic injury model

For inducing the liver injury 7 to 8 week-old male or female mice were given choline-deficient diet (EnvigoTeklad Diets, TD.140207 Choline Deficient Diet with 20% Lard and Irradiated) for one week and then supplemented with 0.10% ethionine in drinking water (Acros Organics,146170100) for another 2 weeks. The mice were either sacrificed at the end of diet or changed to choline-sufficient diet along with normal drinking water until termination.

#### Tamoxifen administration and viral infections

Tamoxifen (Sigma-Aldrich) was dissolved in corn oil. For experiments involving *Krt*19-Cre^ERT2^ x *R26*^LSL^ tdTomato lines, three individual intraperitoneal tamoxifen injections (20mg/ml) were given at a dose of 4mg/20gm body weight of mice for three alternate days. The AAV8 viral vectors were administered 2 weeks after the last tamoxifen injection. p21 plasmid was packaged into the AAV8.TBG vector by the Penn Vector Core. The null vector, AAV8-Ctrl plasmid (AAV8.TBG.PI.Null.bGH, Cat No: AV-8-PV0148) was also obtained from the Penn Vector Core. Viral vectors were diluted in sterile PBS and administered by tail vein or retro-orbital injections with BD Ultra-Fine Insulin Syringes at a viral titer of 5 x 10^11^ copies in total volume of 100 uL. For experiments involving *Kdr*-2A-Cre^ERT2^-2A-eYFP, *R26*^LSL^tdTomato lines, additional 3 tamoxifen injections were given to the mice around the injury as indicated in Figure 4C and Figure 5A.

#### mRNA-LNP formulation and delivery

mRNAs were produced as previously described(Pardi et al., 2013) using T7 RNA polymerase (Megascript, Ambion) on a linearized plasmid encoding codon-optimized (Thess et al., 2015) vascular endothelial growth factor A (VEGFA) (Table S2) or firefly luciferas. mRNAs were transcribed to contain 101 nucleotide-long poly(A) tails. One-methylpseudouridine (m1Ψ)-5’-triphosphate (TriLink) instead of UTP was used to generate modified nucleoside-containing mRNA. RNAs were capped using the m7G capping kit with 2’-O-methyltransferase (ScriptCap, CellScript) to obtain cap1. mRNAs were purified by Fast Protein Liquid Chromatography (FPLC) (Akta Purifier, GE Healthcare) or with cellulose purification as described(Weissman et al., 2013). All mRNAs were analyzed by agarose gel electrophoresis and were stored frozen at −20°C. All nucleoside-modified mRNA are available from the company RNAx created by Dr. Drew Weissman. Poly(C) RNA (Sigma) and purified m1Ψ-containing VEGFA and luciferase mRNAs were encapsulated in LNP using a self-assembly process in which an aqueous solution of mRNA at pH=4.0 is rapidly mixed with a solution of an ionizable cationic lipid, phosphatidylcholine, cholesterol, and polyethylene glycol-lipid dissolved in ethanol(Pardi *et al*., 2015). The ionizable cationic lipid (pKa in the range of 6.0-6.5, proprietary to Acuitas Therapeutics) and LNP composition are described in the patent application WO 2017/004143. They had a diameter of ∼80 nm with a polydispersity index of <0.1 as measured by dynamic light scattering using a Zetasizer Nano ZS (Malvern Instruments Ltd, Malvern, UK) instrument and an encapsulation efficiency of ∼95% as determined using a Ribogreen assay. Acuitas will provide the LNP used in this work to academic investigators who would like to test it. mRNA-LNP formulations were stored at −80°C at a concentration of mRNA of ∼1 μg/μl. VEGFA-mRNA-LNP as well as control Poly(C) RNA-LNP and Luciferase mRNA-LNP were delivered intravenously through retro-orbital sinus. Prior to administration, mRNA-LNPs were thawed and diluted on ice in Dulbecco’s Phosphate Buffered Saline (PBS). Mice were injected with 50 μL of diluted mRNA-LNP (10 μg) intravenously by retro-orbital injections using BD Ultra-Fine Insulin Syringes.

#### Animal tissue harvesting, cryopreservation and serum collection

Mice were sacrificed at their indicated end-points. The liver was separated into its respective lobes, collected directly in 4% paraformaldehyde (PFA), and kept overnight at 4°C prior to OCT embedding. For cryopreservation, tissues were washed thrice with PBS and dipped in 15% sucrose solution for 15 min and transferred to and kept in 30% sucrose solution till they sunk to the bottom. The tissues were then embedded in OCT. The cryopreserved blocks were stored at −80°C. Livers were sectioned at 5 µm thickness using a cryostat (model CM3050 S; Leica, Wetzlar, Germany) and stored at −20°C until required for subsequent staining. For serum separation, blood was collected immediately after euthanizing the mice and prior to liver extraction and kept at room temperature for 30 min. Serum was separated from blood cells by centrifugation at 2500 x g for 15 min.

#### Histology, immunohistochemistry and immunofluorescence image analysis

The frozen sections were allowed to defrost and dry at room temperature for 30 min. The slides were dipped in PBS for 10 min and permeabilized using 0.3% triton X in PBS for 10 min. The slides were rinsed thrice in PBS, 10 minutes each, and blocked with 3% normal donkey serum for 30 min. The sections were then incubated overnight at 4°C with appropriate antibody diluted in PBS. Following primary antibody incubations, the slides were washed thrice with PBS, 10 min each, and incubated with the corresponding fluorescent labeled secondary antibodies for 1h at room temperature protected from light. The slides were finally washed, incubated with Dapi for 3-5 min, rinsed, and mounted using FluorSave reagent (EMD Millipore Corp., 345789). ImageJ version 2.3.0/1.53f was used for image analysis as well as quantifications.

##### Periodic acid–Schiff staining

For staining hepatic glycogen, PAS Staining kit from Sigma Aldrich (395B) was used. Briefly, PFA-fixed, frozen tissue sections were brought to room temperature and hydrated with dH_2_O for 10 min. Subsequently, sections were oxidized in 1% periodic acid for 5 min, rinsed in several changes of dH_2_O and incubated in Schiff’s Reagent for 15 min. After rinsing in dH_2_O twice for 5 min each, the tissue was counterstained with hematoxylin for 15 seconds, rinsed, and dehydrated before mounting with permanent mounting solution.

##### H&E staining

PFA-fixed frozen sections were washed in tap water and dipped for 5 min in hematoxylin (Gill’s or Harris), followed by washing in tap water twice for 2 min. Sections were treated with Bluing Reagent (ammonia water) for 10-15 seconds, washed twice and dipped in 100% ethanol. Eosin was applied to sections for 15 sec, thereafter, the slides were washed several times in 100% ethanol, cleared with Histoclear for 5 min, and mounted using permanent mounting media.

##### Oil Red O staining

Lipid staining was performed on frozen-fixed liver tissue sections using the isopropanol method. Briefly, sections were rinsed with 60% isopropanol and stained with freshly prepared Oil Red O (Sigma) solution for 15 min. Slides were rinsed twice with 60% isopropanol and nuclei were lightly stained using hematoxylin solution. Slides were washed, mounted, and observed under bright-field microscope.

##### LipidSpot staining

To detect lipid accumulation in liver tissue, another method based on fluorescent detection of lipids was performed using LipidSpot™ Lipid Droplet Stains from Biotium (70065-T). Liver sections were defrosted for 30 minutes at room temperature, followed by an immersion in 1x PBS for 10 minutes at room temperature. Sections were then incubated with a mixture of LipidSpot (1:1000) and Dapi (1:3000) diluted in PBS for 10 minutes at room temperature. The slides were washed twice with PBS for 10 minutes and then mounted with FluorSave mounting media (Calbiochem) and imaged on Nikon Eclipse Ni-E upright fluorescent microscope.

##### Trichrome staining

Connective Tissue Staining was performed using Trichrome Staining kit (ab150686, Abcam) following the manufacturer’s instructions. All materials were equilibrated and prepared at room temperature just prior to use. Frozen sections were hydrated in distilled water prior to staining.

#### Serum cholesterol assay

Serum analysis was performed using commercially available Total Cholesterol kit from FUJIFILM Wako Diagnostics (999-02601). Briefly, total cholesterol assay was adapted for a 96-well plate by using 5uL of serum with 200uL of test reagent. After incubation at 37°C for 5 minutes, reactions were read at 600nm and cholesterol levels were calculated from standard curve.

#### Human VEGFA (hVEGFA) ELISA with mouse serum

To detect protein expression from hVEGFA-mRNA, serum levels of hVEGFA were analyzed using VEGFA human ELISA Kit (ab119566, Abcam) following the manufacturer’s instructions. Mice (n=3) were injected with hVEGFA mRNA-LNP (10μg) through retro-orbital sinus injection and their serum was collected 5h, 24h, 48h, and 72h post-injection through submandibular bleed. Serum samples collected at 5h were diluted 10000-fold, 24h samples were diluted 5000-fold, 48h samples were diluted 10-fold while those collected 72h post-injection were diluted 2-fold. Prior to use all reagents were prepared and equilibrated at room temperature. Samples along with the standards were processed according to the manufacturers protocol and the hVEGFA concentrations were calculated from the standard curve.

#### Flow cytometry of isolated non-parenchymal cells and hepatocytes

Liver was perfused according to the previously published method(Everton *et al*., 2021). Hepatocyte and non-parenchymal cell fractions were seeded into 100 μl wells and treated with 1 μg/100 μl Fc Block for 10 minutes at room temperature. Wells were then incubated with primary conjugated antibodies for 20 minutes at room temperature. Following antibody incubation, cells were washed and resuspended in FACS buffer. Dapi (1:100, Invitrogen, R37606) was added to the flow tubes 5 min prior to flow run. Cells were run on BD LSR II SORP flow cytometer and data analyzed with FlowJo v10 software. Care was taken to include appropriate compensation controls for each run. UltraComp eBeads (Invitrogen, 01-2222-42) were used for running compensation controls for APC, APC-Cyanine7 and BV605 conjugated antibodies, while Dapi treated non-parenchymal cell (NPC) fractions from a non-tdTomato mouse were used to compensate Dapi channel. Likewise, unlabeled NPCs from tdTomato mouse were used to compensate for tdTomato channel. At least 1,00,000 cells were analyzed for each fraction. Cells in both the hepatocyte fraction or non-parenchymal cell fraction were first gated to eliminate dead cells positive for Dapi. The SSC and FSC parameters were used to eliminate cell debris. The cells were then gated to exclude blood cells positive for Ter119/CD45/CD11b/CD31. The remaining cells in NPC fraction were gated for EpCAM+ and then evaluated for tdtomato positive cells as shown in the gating strategy, while those in hepatocyte fractions were analyzed for tdtomato (Figure S1B).

#### Experiments on zebrafish

All zebrafish experiments were performed under the approval of the IACUC at the University of Pittsburgh. Embryos and adult fish were raised and maintained under standard laboratory conditions. We used the following transgenic lines: *Tg(fabp10a:CFP-NTR)s931*, *Tg(Tp1:H2B-mCherry)s939*, *Tg(hs:sflt1)bns80*, *Tg(Tp1:CreERT2)s959*, and *Tg(hs:loxP-mCherry-loxP-hVEGFA)mn32*. Their full and official names are listed in resource table.

##### Hepatocyte ablation using the Tg(fabp10a:CFP-NTR) line

Hepatocyte ablation was performed by treating Tg(fabp10a:CFP-NTR) larvae with 10 mM Mtz in egg water supplemented with 0.2% DMSO and 0.2 mM 1-phenyl-2-thiourea from 3.5 to 5 dpf for 36 hours, as previously described (Choi *et al*., 2014).

##### SU5416 treatment

SU5416 (MedChemExpress, Monmouth Junction, NJ) was used at 1.5 µM.

##### Heat-shock conditions for zebrafish larvae

Both Tg(hs:sflt1) and Tg(hs:loxP-mCherry-loxP-hVEGFA) larvae were heat-shocked by transferring them into egg water pre-warmed to 38.5℃ and keeping them at this temperature for 20 minutes, as previously described(Shin et al., 2007).

##### Whole-mount in situ hybridization and immunostaining in zebrafish

Whole-mount in situ hybridization was performed, as previously described(Alexander et al., 1998), with gc(Noël et al., 2010) and f5 probes. For f5 probe synthesis, cDNA from 5-dpf larvae was used as a template together with a forward (5′-CCCTCCTGGCATTCCTGTGTC-3′) and a reverse (5′-<u>TAATACGACTCACTATAGGG</u>CATGGTGGGTCTGCAGCTGT-3′) primer pair to amplify f5; its PCR products were used to make in situ probes. [Underlined is T7 primer sequence.] Whole-mount immunostaining was performed, as previously described(Choi et al., 2015), with mouse anti-Bhmt (1:500; gift from Jinrong Peng at Zhejiang University) and Alexa Fluor 647-conjugated secondary antibodies (1:500; Thermo Fisher Scientific, Waltham, MA).

##### Quantification of Bhmt+ cell number and Bhmt+ area in the zebrafish liver

For the quantification of Bhmt+ cell number, confocal projection images consisting of 6 optical-section images, with 3-μm interval, were used. Bhmt+ and Tp1:H2B-mCherry+ cells were manually counted; the number of Bhmt/Tp1:H2B-mCherry double-positive cells was divided by the number of Tp1:H2B-mCherry+ cells. For the quantification of Bhmt+ liver area, confocal projection images showing Bhmt and fabp10a:CFP-NTR expression were used. Both Bhmt+ and fabp10a:CFP-NTR+ areas in the liver were calculated by ImageJ and the former area was divided by the latter. **Image acquisition, processing, and statistical analysis of zebrafish data:** Zeiss LSM700 confocal and Leica MZ16 microscopes were used to obtain image data. Confocal stacks were analyzed using the Zen 2009 software. All figures, labels, scale bars and outlines were assembled or drawn using the Adobe Illustrator software. Statistical analyses were performed using the GraphPad Prism software. Differences between groups were tested by unpaired Student’s t-tests and considered statistically significant when P < 0.05 (*P < 0.05, **P < 0.01, ***P < 0.001, ****P < 0.0001). Quantitative data were shown as mean ± standard error of the mean (SEM).

##### Human liver tissue staining

Paraffin-embedded liver samples from diseased donors were used as normal liver tissue (n=3). Liver samples from patients with NASH (n=4) and alcoholic cirrhosis (n=1) were fixed overnight in 4% paraformaldehyde (PFA) at 4C. For cryopreservation, tissues were washed thrice with PBS and dipped in 15% sucrose solution for 15 min and transferred to and kept in 30% sucrose solution till they sunk to the bottom. The tissues were then embedded in OCT. The cryopreserved blocks were stored at −80°C. Livers were sectioned at 5 µm thickness using a cryostat (model CM3050 S; Leica, Wetzlar, Germany) and stored at −20°C until required for subsequent staining. For all immunofluorescence staining on human liver tissues, antigen-retrieval was performed in Tris-EDTA buffer ph=9 (Vector Labs) for 20 min at 95C followed by conventional permeabilization, blocking and antibody incubation steps as detailed earlier.

##### Western blot

All samples were incubated with RIPA lysis buffer (Sigma-Aldrich, Saint Louis, MO) containing 1x Halt™ Protease and Phosphatase Inhibitor Cocktail (Thermo Fisher Scientific, Waltham, MA) and incubated for 30 min at 4°C. Samples were centrifuged at 13,000xg for 10 min at 4°C. The supernatant from each sample was then transferred to a new microfuge tube and was used as the whole cell lysate. Protein concentrations were determined by comparison with a known concentration of bovine serum albumin using a Pierce BCA Protein Assay Kit (Thermo Fisher Scientific, Waltham, MA). About 30 µg of lysate were loaded per lane into 10% Mini-PROTEAN TGX gel (BioRad, Hercules, CA). After SDS-PAGE, proteins were transferred onto PVDF membrane (Thermo Fisher Scientific, Waltham, MA). Membranes were incubated with primary antibody solution overnight and then washed. Membranes were incubated for 1 hour in secondary antibody solution and then washed. Target antigens were finally detected using SuperSignal™ West Pico PLUS chemiluminescent substrate (Thermo Fisher Scientific, Waltham, MA). Images were scanned and analyzed using ImageJ software.

##### Real-time PCR

cDNA samples from human cirrhotic and normal hepatocytes were generated from 2ug of RNA using the RevertAid First Strand cDNA Synthesis Kit (Thermo Scientific, #K1621). Commercially available human liver total RNA (Invitrogen, AM7960) was used as positive control. Information regarding primers used in the study are provided in Table S3. Real-time PCR reaction was performed using SYBR Green PCR Master Mix (Applied Biosystem, 4309155) following manufacturers protocol and run on QuantStudio 6 Flex Real-Time PCR System (Applied Biosystem).

#### Statistical analyses

All statistical analyses were performed using GraphPad Prism 9. For comparison between two mean values a 2-tailed Student’s t-test was used to calculate statistical significance. For comparing multiple groups to a reference group one-way ANOVA followed by Tukey’s multi-comparison test was performed. Quantitative data are shown as mean ± standard deviation (SD) and are considered statistically significant when p < 0.05 (*p < 0.05, **p < 0.01, ***p < 0.001, ****p < 0.0001).

## Data and code availability

The authors declare that all data supporting the findings of this study are available within the paper and its supplementary information files.

This paper does not report original code.

Any additional information required to reanalyze the data reported in this paper is available from the lead contact upon request.

## Acknowledgments

This work was supported by the NIH R01DK133404 (VGE and DS) and R01DK101426 (DS). This work was also partially supported by NIH grants DK099257, DK117881, DK119973, TR002383, DK096990, TR003289 and P30DK120531 (Human Synthetic Liver Biology Core) to A.S.-G. We are grateful to Anna Belkina of the BUSM Flow Cytometry Core for technical and analytical assistance, supported by NIH Grant 1UL1TR001430, and Drs. Greg Miller and Marianne James of the CReM, supported by grants R24HL123828 and U01TR001810. Elissa Everton and Anna Smith are supported by the TL1TR001410 award and F31DK127606-01A1.

## Supplemental Information

Document S1: Figures S1 to S6

Table S1: DNA sequences used as template to in vitro transcribe nucleoside modified mRNA.

Table S2: Human tissue samples for histology evaluation, or gene and protein expression analysis.

Table S3: Primer list for qRT-PCR, related to STAR Methods

