## Supplemental data for "VEGFA mRNA-LNP promotes biliary epithelial cell-to-hepatocyte conversion in acute and chronic liver diseases and reverses steatosis and fibrosis"

### Supplemental Information

#### A. Supplemental Figures

**Figure S1. (Supplemental data for Figure 2)**

**(A)** Density plots from flow cytometry of non-parenchymal cell (NPC) fraction isolated from mouse livers. The NPC fraction was gated to exclude dead cell, debris, and cells expressing CD11b, CD45, Ter119, CD31. Values show % tdTomato+ EpCAM+ population. Lower panel shows cells from a control non-tdTomato mouse run simultaneously with each experimental mouse.

**(B)** Gating strategies of hepatocyte and non-parenchymal fractions.

**(C)** Representative immunofluorescent microscopy images showing extent of ductular response (KRT7+ BECs) in *Krt19*-Cre$^{ERT}$, *R26*$^{LSL}$tdTomato mice treated with AAV8-Tbg-p21 vector and choline-deficient diet with 0.1% ethionine supplementation for 2 weeks, followed by VEGFA mRNA-LNP or control Poly(C) RNA-LNP injections, 10 ug/20g body weight. The bar graph shows quantification of KRT7-positive area averaged from at least three different fields in each mouse in the two groups (n=4 per group).

**(D)** KRT7 and tdTomato staining showing decrease in BEC density in regions of BEC-to-hepatocyte conversion in VEGFA mRNA-LNP-treated group (outline).

**(E)** Scheme showing key interventions in the experimental design using the *Krt19*-Cre$^{ERT}$, *R26*$^{LSL}$tdTomato mice without inducing hepatocyte senescence through AAV8-Tbg-p21 vector. Injury was induced by choline-deficient diet with 0.1% ethionine supplementation for 2 weeks, followed by VEGFA mRNA-LNP or control Poly(C) RNA-LNP injections, 10 ug/20g body weight, as indicated.

**(F)** Representative immunofluorescent microscopy images showing extent of ductular response (KRT7+ biliary epithelial cells) in chronically injured liver in the absence of AAV8 p21-induced hepatic senescence.

**(G)** Close-up images of tdTomato+ liver cells in Poly(C) RNA-LNP- and VEGFA mRNA-LNP-treated mice.

**(H)** Representative immunofluorescent microscopy images showing tdTomato+ liver cells and p21 staining (green) on serial sections of liver tissue demonstrating the endogenous up-regulation of p21 in areas around tdTomato+ hepatocytes. Images on the right have been magnified to reveal p21 staining.

Numerical data are presented as mean ± s.d. P values were determined by one-way ANOVA followed by Tukey's method for multiple comparisons. $^{**}p < 0.01$, $^{***}p < 0.001$. ns = non-significant.

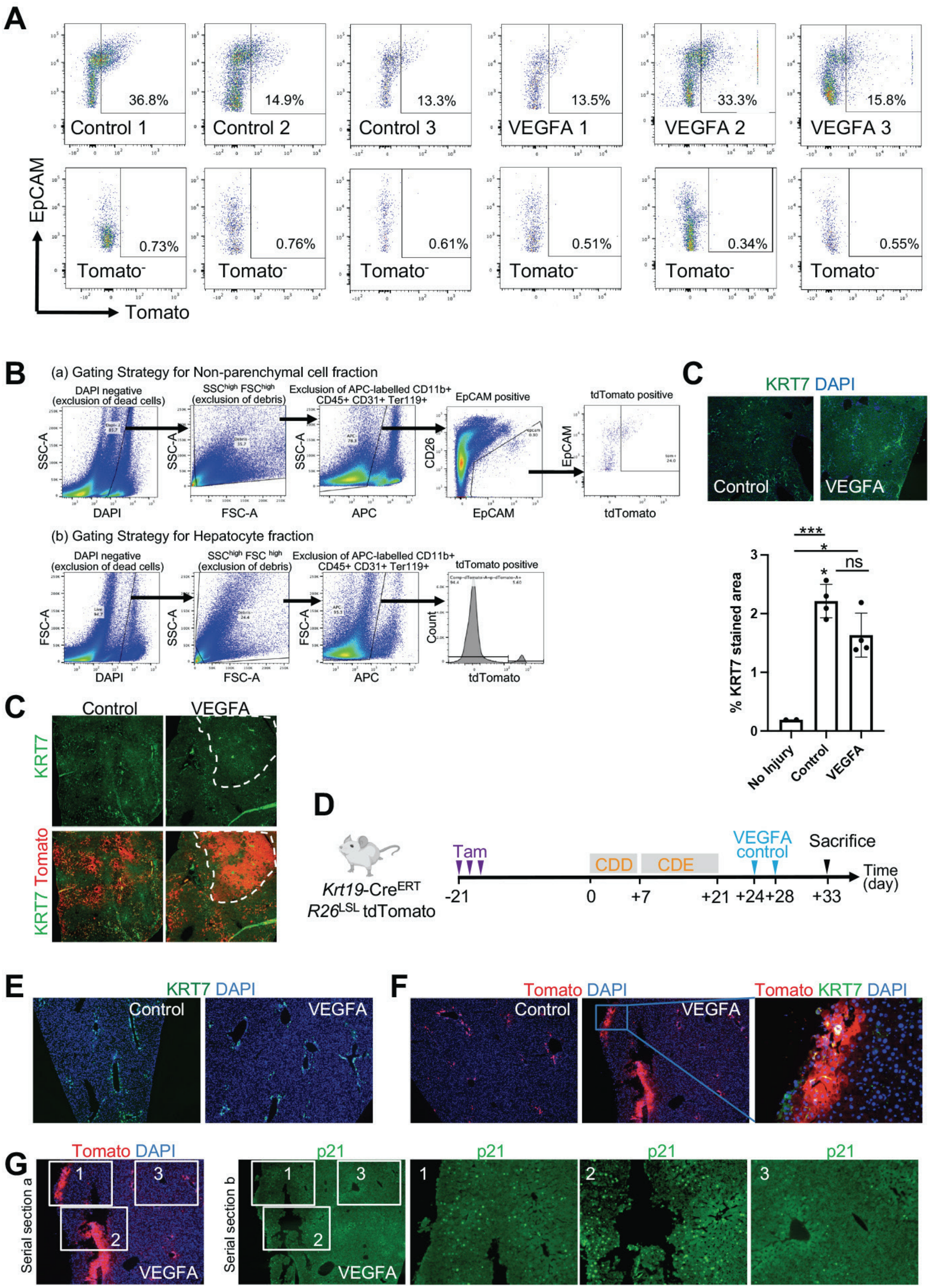
A
36.8%
Control 1
14.9%
Control 2
13.3%
Control 3
13.5%
VEGFA 1
33.3%
VEGFA 2
15.8%
VEGFA 3
EpCAM
0.73%
0.76%
0.61%
0.51%
0.34%
0.55%
Tomato⁻
Tomato
B
(a) Gating Strategy for Non-parenchymal cell fraction
DAPI negative (exclusion of dead cells)
SSC^high FSC^high (exclusion of debris)
Exclusion of APC-labelled CD11b+ CD45+ CD31+ Ter119+
EpCAM positive
tdTomato positive
SSC-A
DAPI
FSC-A
APC
CD26
EpCAM
tdTomato
(b) Gating Strategy for Hepatocyte fraction
DAPI negative (exclusion of dead cells)
SSC^high FSC ^high (exclusion of debris)
Exclusion of APC-labelled CD11b+ CD45+ CD31+ Ter119+
tdTomato positive
FSC-A
SSC-A
Count
C
KRT7 DAPI
Control
VEGFA
% KRT7 stained area
***
*
ns
No Injury
Control
VEGFA
C
Control
VEGFA
KRT7
KRT7 Tomato
D
Krt19-Cre^ERT
R26^LSL tdTomato
Tam
CDD
CDE
VEGFA
control
Sacrifice
Time (day)
-21
0
+7
+21
+24
+28
+33
E
KRT7 DAPI
Control
VEGFA
F
Tomato DAPI
Control
VEGFA
Tomato KRT7 DAPI
G
Tomato DAPI
Serial section a
VEGFA
p21
Serial section b
1
2
3

**Figure S2. (Supplemental data for Figure 3)**

**(A)** Representative immunofluorescence microscope images showing expression of p21-induced by AAV8-Tbg-p21 vector as compared to mice injected with a null vector. Mice were administered with $5x10^{11}$ gc of AAV8-Tbg-p21 vector or null vector intravenously through retro-orbital sinus injection.

**(B)** Density plots from flow cytometry of NPC fraction isolated from mouse livers. The NPC fraction was gated to exclude dead cell, debris, and cells expressing CD11b, CD45, Ter119, CD31. Values show % tdTomato+ EpCAM+ population. Lower panel shows cells from a control non-tdTomato mouse run simultaneously with each experimental mouse.

**(C)** Scheme showing key interventions in the experimental design using the *Krt19*-$Cre^{ERT}$, *R26*$^{LSL}$tdTomato mice without administering AAV8-Tbg-p21 vector to induce hepatocyte senescence. Injury was induced by a single intraperitoneal injection of APAP (500mg/kg to female mice) followed by VEGFA mRNA-LNP or control Poly(C) RNA-LNP injections, 10 $\mu$g/20g body weight.

**(D)** Representative close-up images of tdTomato+ liver cells in Poly(C) RNA-LNP- and VEGFA mRNA-LNP-treated mice. HNF4$\alpha$ (green) staining shows the hepatocyte identity of selected tdTomato+ cluster.

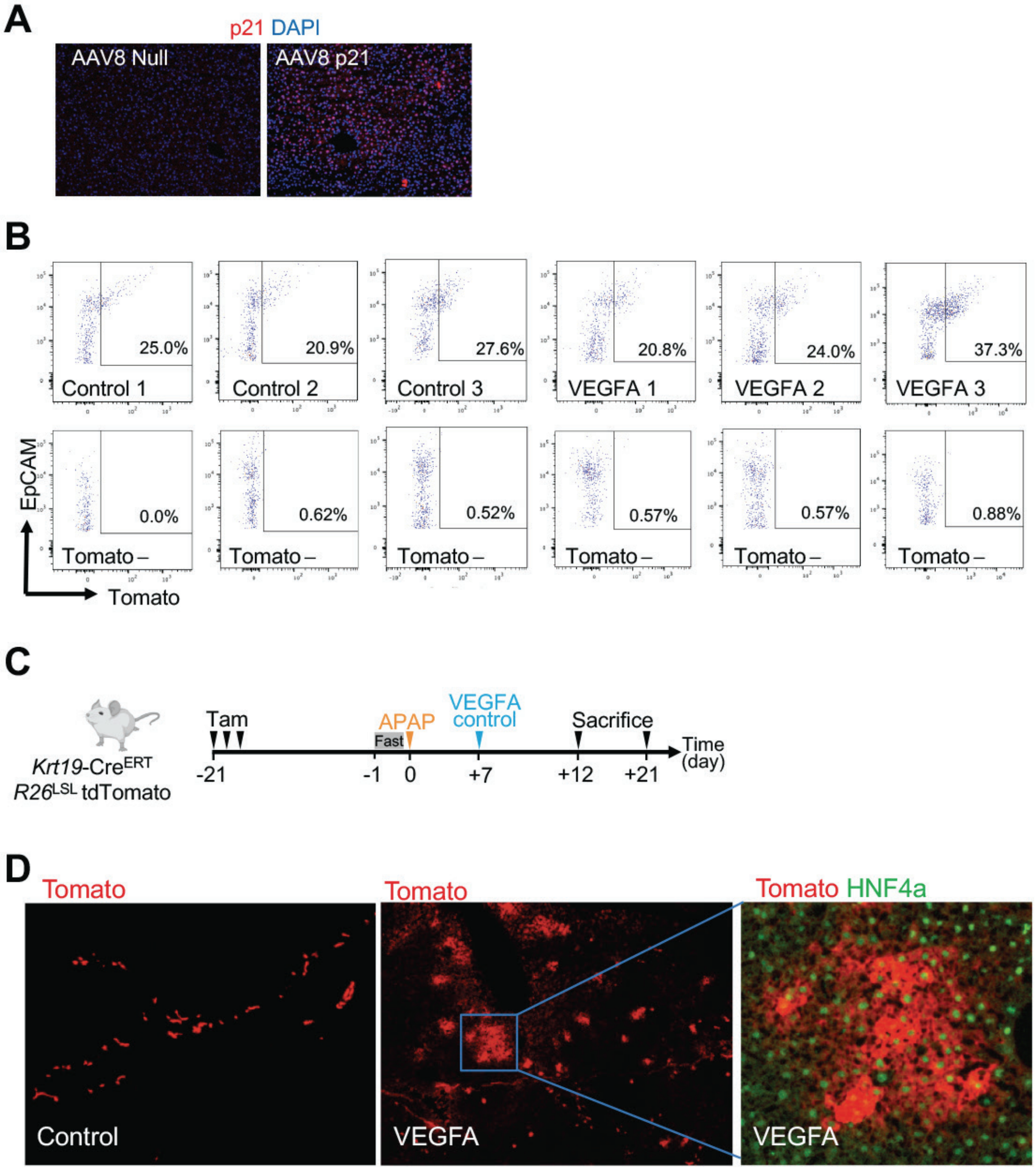
A
p21 DAPI
AAV8 Null
AAV8 p21
B
25.0%
Control 1
20.9%
Control 2
27.6%
Control 3
20.8%
VEGFA 1
24.0%
VEGFA 2
37.3%
VEGFA 3
EpCAM
Tomato
0.0%
Tomato –
0.62%
Tomato –
0.52%
Tomato –
0.57%
Tomato –
0.57%
Tomato –
0.88%
Tomato –
C
Krt19-Cre^ERT
R26^LSL tdTomato
Tam
Fast
APAP
VEGFA
control
Sacrifice
Time
(day)
-21
-1
0
+7
+12
+21
D
Tomato
Control
Tomato
VEGFA
Tomato HNF4a
VEGFA

**Figure S3. (Supplemental data for Figure 4)**

**(A)** Immunofluorescent images showing co-expression of the endothelial cell markers KDR and CD31 in the liver of uninjured mice.

**(B)** Schematic structure of the *Kdr*-2A-$Cre^{ERT2}$-2A-eYFP construct.

**(C)** Representative immunofluorescent images showing specificity and sensitivity of the *Kdr*-2A-$Cre^{ERT2}$-2A-eYFP construct in *Kdr*-2A-$Cre^{ERT2}$-2A-eYFP mice. YFP (green) expression overlaps the KDR+ (red) expression in the liver endothelial cells.

**(D)** Scheme showing the experimental design for defining leakiness of Cre recombinase in *Kdr*-2A-$Cre^{ERT2}$-2A-eYFP, $R26^{LSL}$tdTomato mice. Mice were administered with three corn oil injections on alternate days and sacrificed 10 days later.

**(E)** Representative immunofluorescent microscope images showing leakiness of Cre in *Kdr*-2A-$Cre^{ERT2}$-2A-eYFP, $R26^{LSL}$tdTomato mice (male, n=3; female n=3). Enlarged images of selected areas demonstrate KDR+ endothelial cell identity (green) of tdTomato+ cells.

**(F)** Representative immunofluorescent microscope images of KDR+ and tdTomato+ cells and their corresponding pixel representation by ImageJ used for quantitation of percent leakiness. The bar graph depicts quantification of % tdTomato+ KDR+ cells averaged from at least four different fields in each mouse in the two groups (n=3 per group).

**(G)** Bar graph representing quantitation of rare tdTomato+ hepatocytes observed in *Kdr*-2A-$Cre^{ERT2}$-2A-eYFP, $R26^{LSL}$tdTomato mice treated with corn oil. Numerical data are presented as mean ± s.d. P values were determined by two-tailed Student’s t-test. ns=non-significant.

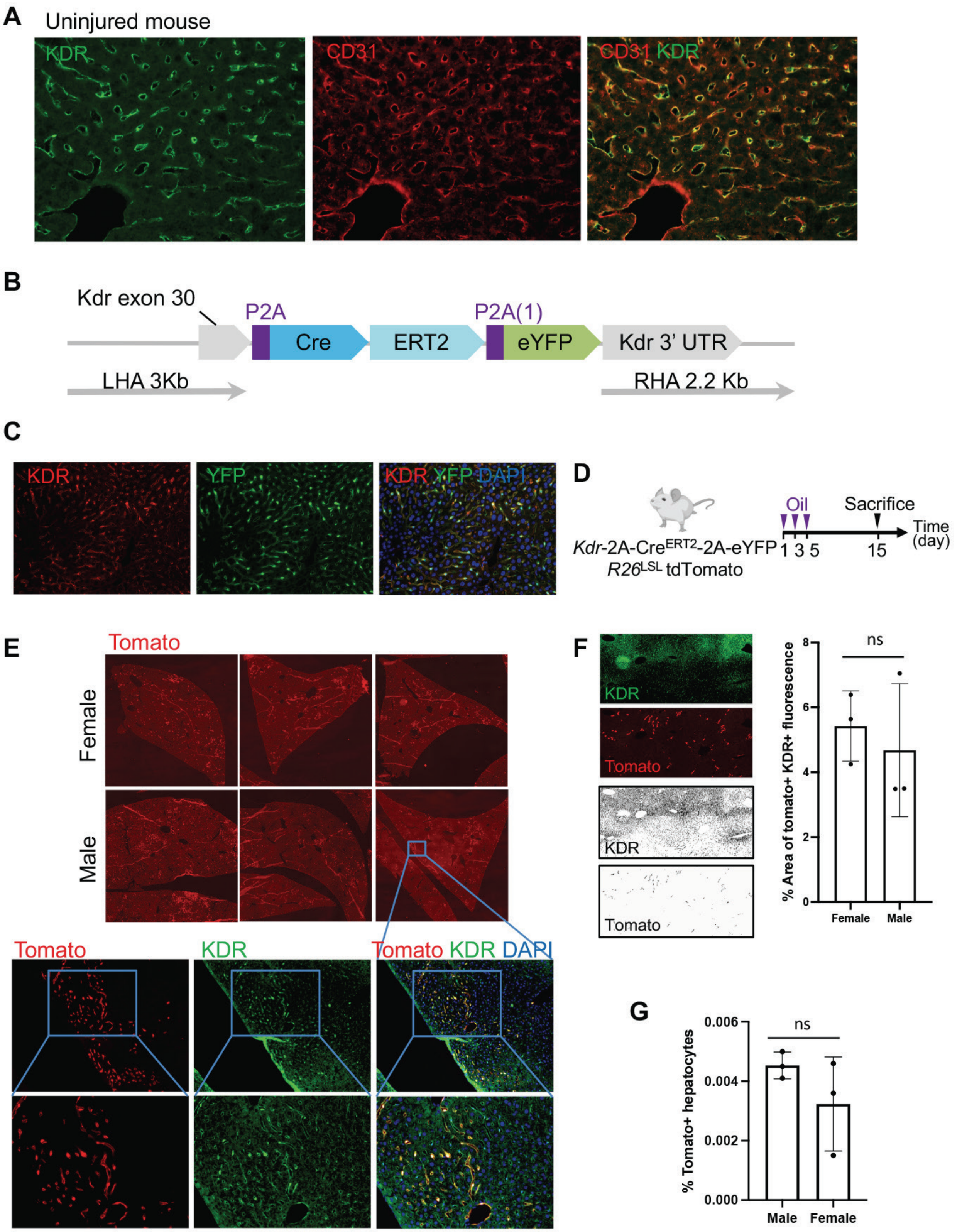
A
Uninjured mouse
KDR
CD31
CD31 KDR
B
Kdr exon 30
P2A
Cre
ERT2
P2A(1)
eYFP
Kdr 3' UTR
LHA 3Kb
RHA 2.2 Kb
C
KDR
YFP
KDR YFP DAPI
D
Oil
Sacrifice
Time (day)
1 3 5
15
Kdr-2A-CreERT2-2A-eYFP
R26LSL tdTomato
E
Tomato
Female
Male
Tomato
KDR
Tomato KDR DAPI
F
KDR
Tomato
KDR
Tomato
ns
% Area of tomato+ KDR+ fluorescence
Female
Male
G
ns
% Tomato+ hepatocytes
Male
Female

**Figure S4. (Supplemental data for Figure 4 and 5)**
**(A)** Representative image showing lineage tracing efficiency in KDR expressing cells. The labelling was uniform and complete in endothelial cells across all mice as shown.
**(B)** Representative immunofluorescent microscope images showing ductular response (EpCAM+ BECs) in *Kdr*-2A-Cre$^{ERT2}$-2A-eYFP, *R26*$^{LSL}$tdTomato mice treated with AAV8-Tbg-p21 vector and choline-deficient diet with 0.1% ethionine supplementation for 2 weeks, followed by VEGFA mRNA-LNP or control firefly luciferase mRNA-LNP injections, 10 μg/20g body weight. The bar graph shows quantification of area of EpCAM+ staining averaged from at least three different fields in each mouse in the two groups (n=3 per group).
**(C)** Krt7 staining showing comparison of ductular response in *Kdr*-2A-Cre$^{ERT2}$-2A-eYFP, *R26*$^{LSL}$tdTomato mice treated with AAV8-Tbg-p21 vector and APAP (500mg/kg to female mice, 300mg/kg to male mice) followed by VEGFA mRNA-LNP or control Luciferase mRNA-LNP injections, 10 μg/20g body weight. The bar graph represents quantification of Krt7 stained area averaged from at least three different fields in each mouse in the two groups (n=6 per group).
**(D)** Scheme showing the experimental design using the *Kdr*-2A-Cre$^{ERT2}$-2A-eYFP, mice to capture YFP (or KDR) expressing cells after acute liver toxicity. AAV8-Tbg-p21 vector was administered to induce hepatocyte senescence and injury was induced by a single intra-peritoneal injection of acetaminophen (400mg/kg APAP to male mice and 500mg/kg to female mice). Mice were sacrificed three days after the injury.
Numerical data are presented as mean ± s.d. P values were determined by one-way ANOVA followed by Tukey's method for multiple comparisons. ***p< 0.001, ****p< 0.0001, ns=non-significant.

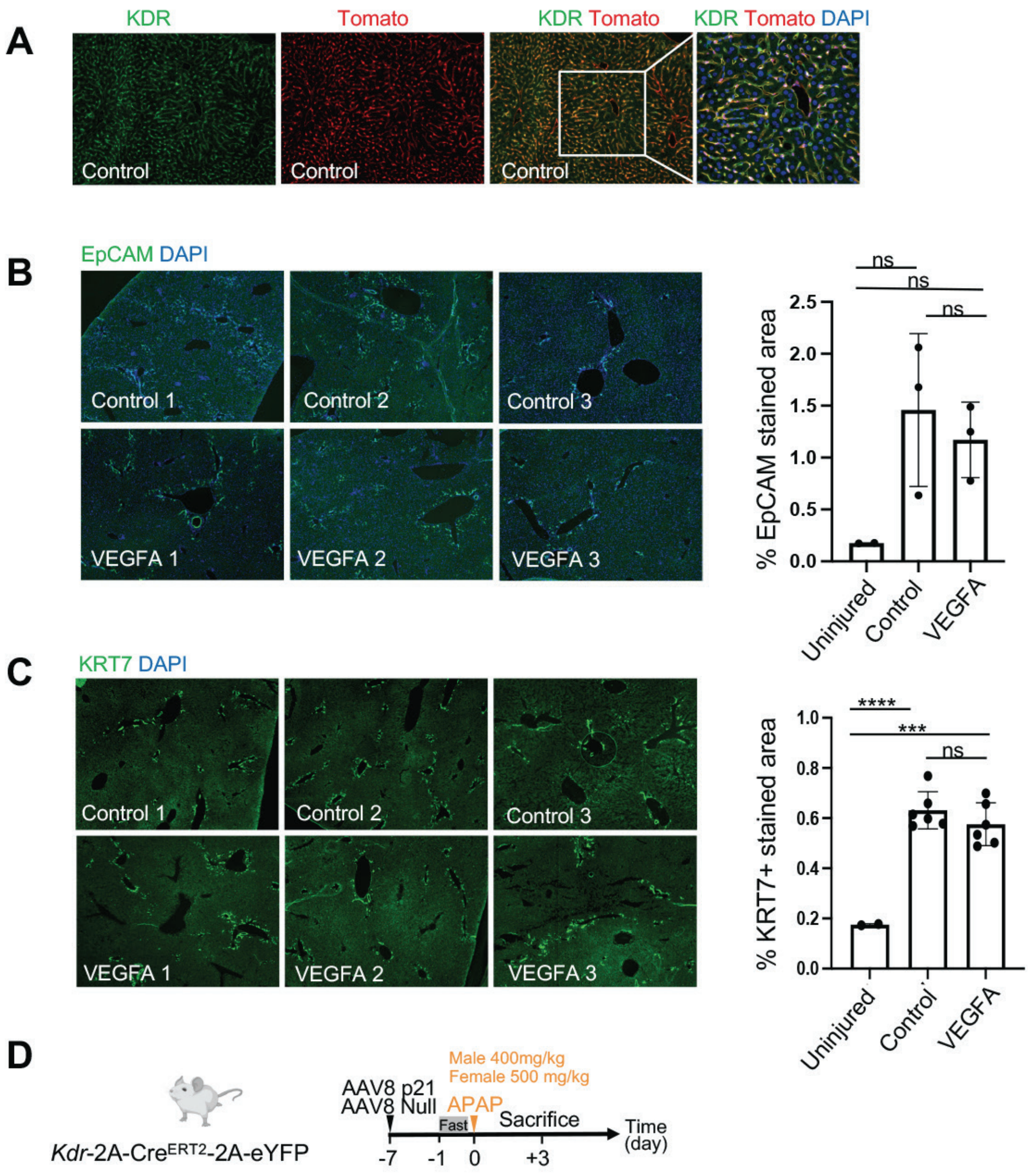
A
KDR
Tomato
KDR Tomato
KDR Tomato DAPI
Control
Control
Control
B
EpCAM DAPI
Control 1
Control 2
Control 3
VEGFA 1
VEGFA 2
VEGFA 3
% EpCAM stained area
ns
ns
ns
Uninjured
Control
VEGFA
C
KRT7 DAPI
Control 1
Control 2
Control 3
VEGFA 1
VEGFA 2
VEGFA 3
% KRT7+ stained area
****
***
ns
Uninjured
Control
VEGFA
D
Kdr-2A-Cre^ERT2-2A-eYFP
AAV8 p21
AAV8 Null
Male 400mg/kg
Female 500 mg/kg
APAP
Fast
Sacrifice
Time
(day)
-7
-1
0
+3

**Figure S5. (Supplemental data for Figure 6)**
**(A)** Representative brightfield images of H&E-stained cirrhotic human liver specimens showing the abnormal liver morphology and thick fibrous septa around the portal tracts of the diseased human livers.
**(B)** Brightfield images of trichrome stained human cirrhotic liver tissues showing collagen deposition (blue).
**(C)** Oil Red O-stained human cirrhotic liver tissues showing lipid accumulation in the liver (red).
**(D)** Representative immunofluorescence images showing ductular response (KRT7+ BECs) in the tissues of human liver samples at 4X magnification. The selected regions are shown at 10X magnification to clearly visualize KRT7-positive BECs (red) in the dense fibrous septa as depicted by DAPI staining (blue).

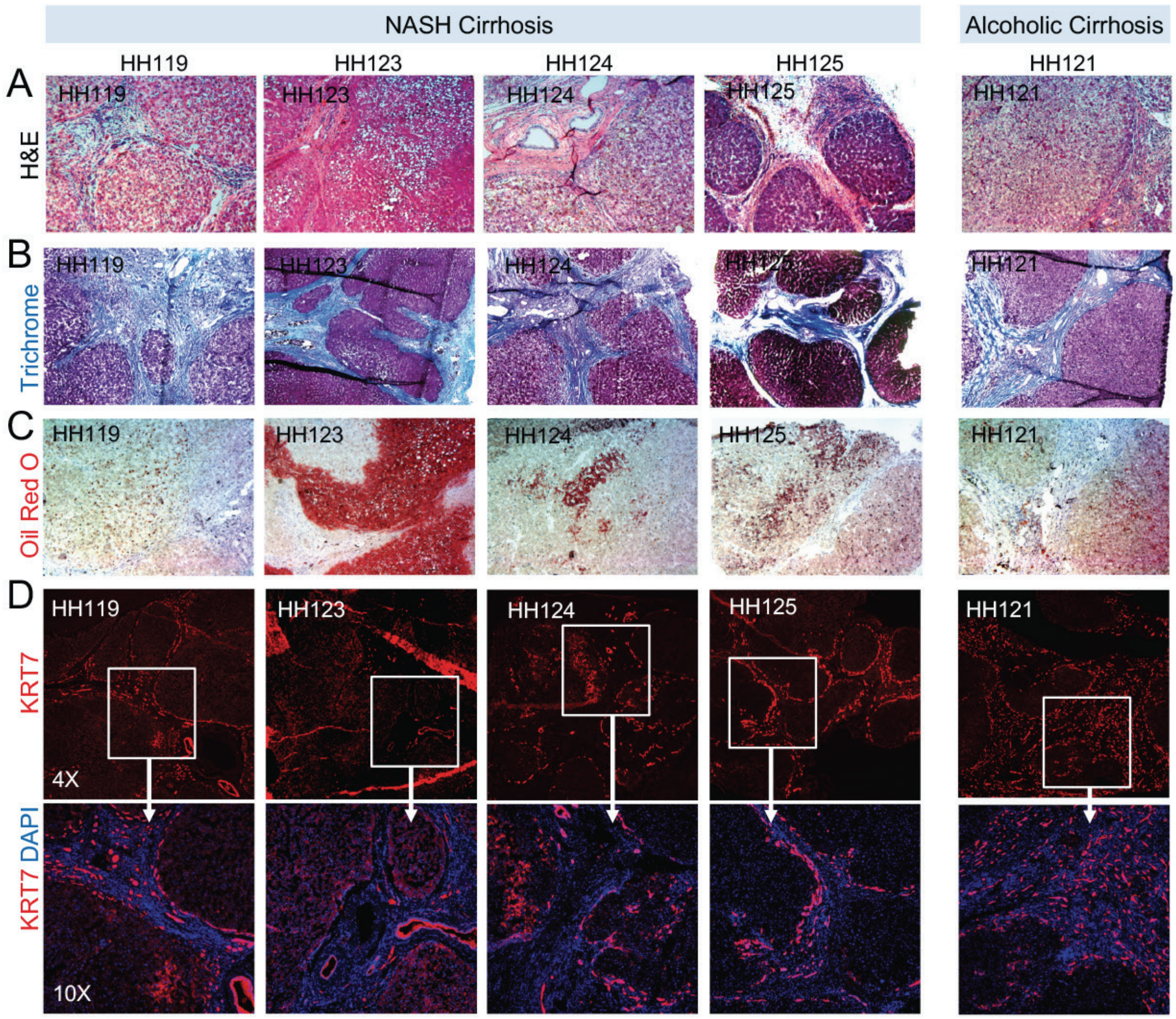

**Figure S6. (Supplemental data for Figure 6)**
**(A)** Representative immunofluorescent images of cirrhotic human liver tissues showing the GS expression (green) in the central vein area away from the fibrous where KRT7+ BECs are located (red).
**(B)** Representative immunofluorescent images of normal human liver tissues (n=3) showing the GS expression (green) in the central vein areas. KRT7 (red) depicts the portal vein area.
**(C)** KDR (red) and KRT7 (green) staining showing the normal endothelial staining pattern in the diseased donor liver tissues.
**(D)** GS and its corresponding IgG isotype staining on serial sections demonstrating the specificity of GS antibody.
**(E)** Relative gene expression of albumin determined in hepatocytes isolated from normal (n=5) or cirrhotic human livers, Child Pugh B (n=5) and Child-Pugh C (n=4).

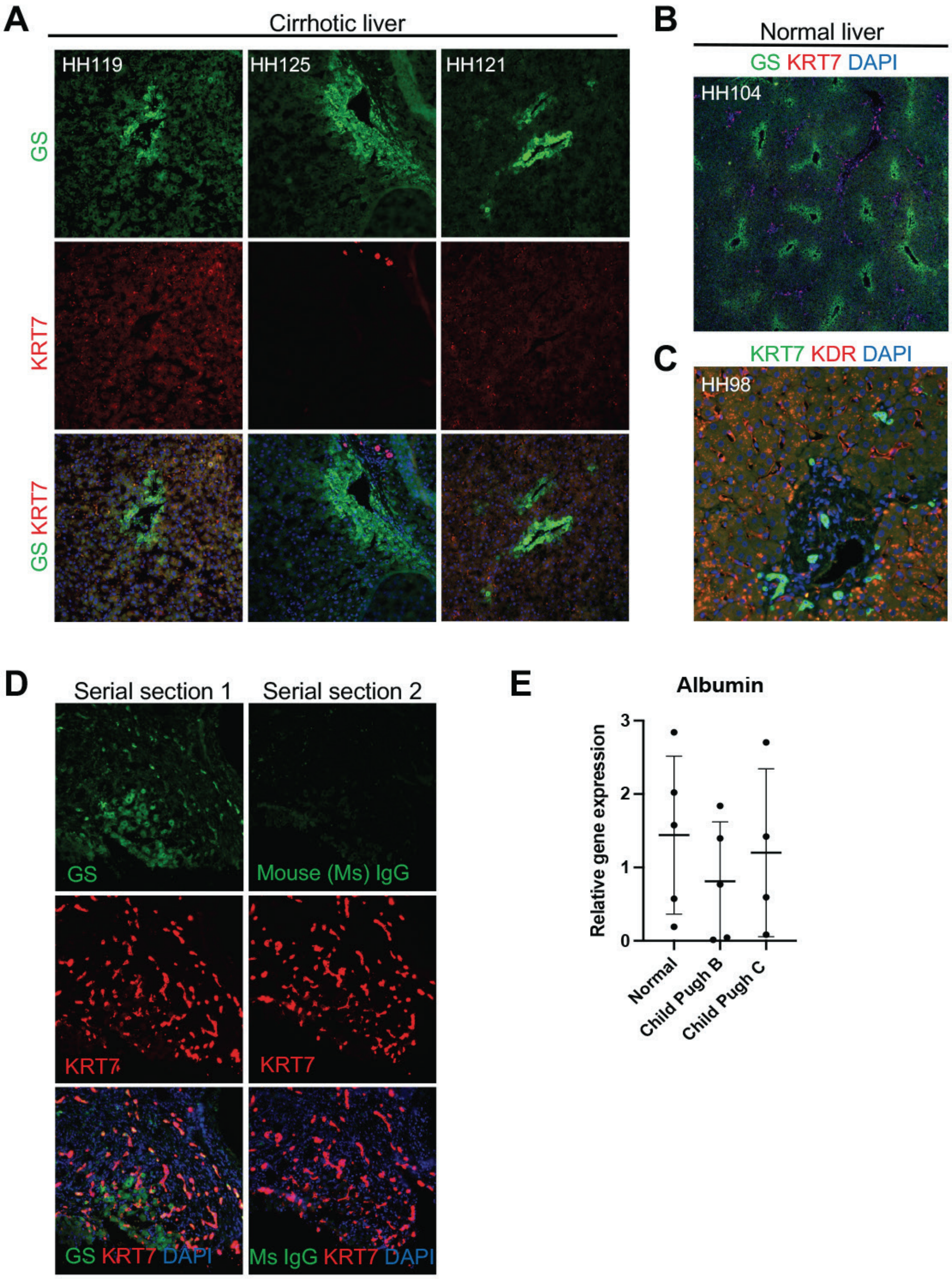
A
Cirrhotic liver
HH119
HH125
HH121
GS
KRT7
GS KRT7
B
Normal liver
GS KRT7 DAPI
HH104
C
KRT7 KDR DAPI
HH98
D
Serial section 1
Serial section 2
GS
Mouse (Ms) IgG
KRT7
KRT7
GS KRT7 DAPI
Ms IgG KRT7 DAPI
E
Albumin
Relative gene expression
0
1
2
3
Normal
Child Pugh B
Child Pugh C

### B. Supplemental Tables

**Table S1:** Details of human tissue samples for histology evaluation, or gene and protein expression analysis.

| Code | Age | Sex | Diagnosis | Child-Pugh Score |
|---|---|---|---|---|
| HH98 | 15 years | male | Donor liver | NA |
| HH104 | 11 years | male | Donor liver | NA |
| HH09 | 37 years | female | FNH/No chemo | NA |
| HH119 | 59 years | male | NASH cirrhosis | B |
| HH123 | 58 years | male | NASH cirrhosis | C |
| HH124 | 63 years | male | NASH cirrhosis | B |
| HH125 | 66 years | male | NASH cirrhosis | B |
| HH121 | 71 years | male | Alcoholic cirrhosis | C |
| HH48 | 33 years | female | HCA “normal” | NA |
| HH55 | 61 years | male | ICCA “normal” | NA |
| HH16 | 44 years | male | Colon Ca “normal” | NA |
| HH36 | 47 years | female | EHE “normal” | NA |
| HH98 | 15 years | male | Donor liver “normal” | NA |
| HH32 | 71 years | female | NASH cirrhosis | B |
| HH72 | 48 years | male | Alcoholic cirrhosis | B |
| HH49 | 50 years | female | NASH cirrhosis | B |
| HH62 | 67 years | male | NASH cirrhosis | B |
| HH22 | 53 years | male | NASH cirrhosis | B |
| HH26 | 52 years | male | Alcoholic cirrhosis | C |
| HH29 | 68yo | female | NASH cirrhosis | C |
| HH97 | 62yo | female | NASH cirrhosis | C |
| HH123 | 58yo | male | NASH cirrhosis | C |

**Table S2:** DNA sequences used as template to in vitro transcribe nucleoside modified mRNA.

| Human VEGF165 |
|---|
| ATGAACTTCCTGCTGTCCTGGGTGCACTGGTCCCTGGCCCTGCTGCTGTACCTGCACCA<br>CGCCAAGTGGTCCCAGGCCGCCCCCATGGCCGAGGGCGGCGGCCAGAACCACCACGA<br>GGTGGTGAAGTTCATGGACGTGTACCAGCGgTCCTACTGCCACCCCATCGAGACCCTGG<br>TGGACATCTTCCAGGAGTACCCCGACGAGATCGAGTACATCTTCAAGCCCTCCTGCGTG<br>CCCCTGATGCGCTGCGGCGGCTGCTGCAACGACGAGGGCCTGGAGTGCGTGCCCACC<br>GAGGAGTCCAACATCACCATGCAGATCATGCGCATCAAGCCCCACCAGGGCCAGCACA<br>TCGGCGAGATGTCCTTCCTGCAGCACAACAAGTGCGAGTGCCGCCCCAAGAAGGACCG<br>CGCCCGCCAGGAGAACCCCTGCGGCCCCTGCTCCGAGCGCCGCAAGCACCTGTTCGT<br>GCAGGACCCCCAGACCTGCAAGTGCTCCTGCAAGAACACCGACTCCCGCTGCAAGGCC<br>CGCCAGCTGGAGCTGAACGAGCGCACCTGCCGCTGCGACAAGCCCCGCCGCTAA |

**Table S3:** List of human primers used for gene expression analysis in isolated human hepatocytes.

| Gene | Primer Sequence | Amplicon length |
|---|---|---|
| hKRT7 | F: GGACATCGAGATCGCCACCT<br>R: ACCGCCACTGCTACTGCCA | 124 |
| h18s rRNA | F: CTCAACACGGGAAACCTCAC<br>R: CGCTCCACCAACTAAGAACG | 110 |
| hKDR | F: ACTTTGGAAGACAGAACCAAATTATCTC<br>R: TGGGCACCATTCCACCA | 52 |
| hAlbumin | F: GTGAAACACAAGCCCAAGGCAACA<br>R: TCCTCGGCAAAGCAGGTCTC | 116 |
| hHNF4a | F: GGTGTCCATACGCATCCTTGAC<br>R: AGCCGCTTGATCTTCCCTGGAT | 144 |
